# Next-Generation Neural Mass Models Reproduce Features of Speech Processing

**DOI:** 10.1101/2025.10.20.683434

**Authors:** Andrew Shannon, David Barton, Martin Homer, Conor Houghton

**Affiliations:** School of Computer Science, University of Bristol, Bristol BS8 1TH, UK; School of Engineering Mathematics and Technology, University of Bristol, Bristol BS8 1TH, UK

## Abstract

Segregation of speech into syllables is a key step in neural speech processing. It relies on the alignment of neural activity with the rhythmic structure of speech. Two competing hypotheses explain this ‘neural speech tracking’, phase-resetting and evoked responses. While phenomenological modelling of these hypotheses has been successful, we still lack understanding of the underlying cortical circuits. To investigate these mechanisms, we evaluate whether a biophysical next-generation neural mass model can reproduce several features of neural speech tracking, using phenomenological models of the competing hypotheses as algorithmic baselines. We investigate the models’ dynamics with four tests: recreating in-silico an EEG experiment that identified a correlation between tracking strength and phoneme sharpness, computing the Phase Concentration Metric, testing the effect of varying syllabic rates, and evaluating the Inter Event Phase Coherence across phoneme onsets. While all of the models that we study reproduce the sharpness-tuned rhythmic speech tracking, the evoked model requires a pre-processed acoustic edge impulse stimulus. We demonstrate that the neural mass model is performing thresholded phase-resetting triggered by sharp onsets in the continuous speech envelope. This produces cross-frequency nested oscillations that qualitatively match an experimentally-observed dual-peak signature in the Inter Event Phase Coherence. Our results indicate that the biophysical neural mass model provides a mechanistic bridge between generic oscillatory dynamics in cortical populations and the cognitive computations of speech tracking. Indeed, the non-linear dynamics of the neural mass model offer an explanation for how peak-rate event representations in auditory cortex activity arise in response to continuous acoustic input.

**Significance Statement:** Syllable segregation is crucial but challenging as natural speech lacks clear boundaries, yet humans perform this computation effortlessly. Speech aligns neural activity to syllabic rhythms, predicting syllable timing, but the underlying cortical mechanisms remain unknown. Relating this macroscopic behaviour to neurobiology is challenging; however, next-generation neural mass models promise to resolve this. We demonstrate that these models reproduce sharpness-tuned tracking and acoustic edge extraction. Dynamical analyses indicate this occurs through thresholded phase-resetting to phoneme onsets, triggering cross-frequency nested oscillations. Our results both advance biophysical understanding of syllable segregation and validate the models’ capacity for simulating macroscopic neural activity. These models offer a bridge between the neurobiology of the auditory cortex and speech processing dynamics that phenomenological models cannot provide.

## Introduction

Humans are extremely good at splitting speech sound into syllables. However, it is not a straightforward task because syllables often arrive in a continuous stream without discrete gaps. In this study, we focus on a key mechanism of syllable segregation: the alignment of neural activity with the syllabic rhythm (Räsänen et al., 2018). This ‘neural speech tracking’ has been causally linked to speech comprehension in stimulation studies (Riecke et al., 2018; Zoefel et al., 2018a).

Multiple mechanisms have been proposed for speech tracking and it is unknown whether tracking is the result of activity evoked by acoustic features or phase-resetting of endogenous oscillations. This is an example of a broader debate in sensory neuroscience (Capilla et al., 2011; Zoefel et al., 2018b; Lalor and Nidiffer, 2025). The evoked-response hypothesis suggests that neural activity tracks speech by responding to discrete features of the envelope such as syllable onsets. Representations of these features have been identified within the superior temporal gyrus (STG) (Hertrich et al., 2012; Oganian and Chang, 2019), and mathematical modelling suggests that evoked-responses can account for the response to natural speech seen in macroscale neural recordings (Zou et al., 2021; Oganian et al., 2023).

The phase-resetting hypothesis, in contrast, assumes that pre-existing oscillatory activity within the auditory cortex is shifted by a stimulus, causing entrainment to the stimulus rhythm. Neuronal oscillations have been implicated in various related functions (Schroeder and Lakatos, 2009; Obleser and Kayser, 2019; Poeppel and Assaneo, 2020; Peelle and Davis, 2012), and they may facilitate speech processing through a hierarchical process using cross-frequency coupling to gate neuronal spiking in time with quasi-rhythmic stimuli (Giraud and Poeppel, 2012).

Both of these mechanisms have primarily been investigated using phenomenological models (Oganian et al., 2023; Doelling et al., 2019; Doelling and Assaneo, 2021). Therefore, although we have an idea of the fundamental dynamics responsible for syllable segregation, they are disconnected from the biological structures that support them. To achieve a deeper understanding, we need models that bridge between the high-level processing of syllable segregation and the cortical circuits enacting it. Biophysical models attempt to provide this connection by incorporating known biological details.

To address this, we explore whether modern biophysical neural mass models are a suitable tool for investigating the mechanistic underpinnings of neural speech tracking. Specifically, the Byrne-Coombes next-generation Neural Mass Model (NMM) (Byrne et al., 2017; Pietras et al., 2019; Byrne et al., 2020, 2022) is designed to model meso-scale oscillatory circuits. We investigate whether the NMM can reproduce established features of neural speech tracking. To help interpret the NMM’s behaviour, we compare its dynamics to phenomenological models of the phase-resetting and evoked-response hypotheses.

As the initial benchmark in our study, we use data from Cucu et al. (2022) showing that neural speech tracking is correlated with phoneme sharpness. This result is in agreement with similar findings in the response to non-speech tones (Oganian and Chang, 2019) and spoken digits (Doelling et al., 2014). Further, it has been identified as a primary driver of cortical speech responses Daube et al. (2019), and shown to normalise to the contextual speech rate (Oganian et al., 2023). This benchmark requires the capture of both gross syllabic-rate tracking and the lower-level feature of sharpness-specific tuning.

Here, we recreate the EEG experiment in-silico, demonstrating that all three models can reproduce the sharpness-specific tuning of neural speech tracking. While this confirms the NMM’s capacity to track speech, it indicates the benchmark is limited in its ability to distinguish between viable mechanisms. As such, we investigate the models’ dynamics further using the Phase Concentration Metric (PCM) (Doelling et al., 2019), a syllabic rate variation test, and Inter Event Phase Coherence (IEPC) analysis. We demonstrate that the NMM tracks the syllabic rate through thresholded phase-resetting and cross-frequency coupling. This is produced by phoneme onsets triggering phase-locked high-frequency oscillations, temporally nested within the slower syllabic rate. This behaviour qualitatively matches the double peaked IEPC feature previously only recreated with linear evoked models (Oganian et al., 2023) and demonstrates that non-linear processing of the continuous speech envelope is sufficient to explain the presence of peak-rate event representations in the STG.

## Materials and Methods

### Models

#### Neural mass model

The NMM, (Byrne et al., 2017), is an exact mean field reduction of an infinite all-to-all connected network of spiking quadratic integrate and fire neurons. It provides a biophyiscally grounded description of the activity of a large population of cortical neurons with parameters corresponding to biophysical features such as membrane and synaptic time constants, synaptic reversal potentials, and the distribution of neuron excitabilities. As the model is based on a spiking neural network, it incorporates spike timing and subsequently captures the synchronisation of the population rather than just the mean firing rate. We used the first variant (Byrne et al., 2017), incorporating *α*-function synaptic filtering of spikes, but not gap junctions or competing excitatory-inhibitory populations (Pietras et al., 2019; Byrne et al., 2020, 2022). We consider only a single population here, representing a macroscopic ensemble of auditory cortical neurons.

The model comprises a single complex differential equation for a Kuramoto order parameter *Z*(*t*), and two input filters of the synaptic conductance *g*(*t*) and external stimulus current *A*(*t*). The Kuramoto order parameter, *Z*(*t*), describes the synchrony of the population’s spikes and satisfies

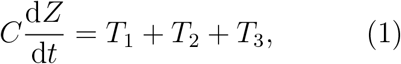

where *C* is the membrane capacitance.

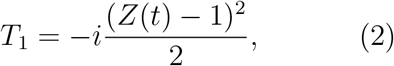

captures the intrinsic, self referential dynamics of the network activity.

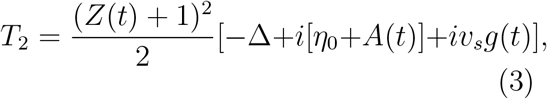

describes the response to driving forces: the internal excitability *η*_0_ and its heterogeneity Δ, an external drive *A*(*t*), and the global post synaptic current *v*_*s*_*g*(*t*) with *v*_*s*_ the synaptic reversal potential.

The third term, in (1),

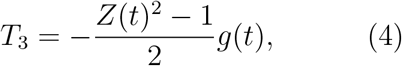

describes the feedback from state dependent synaptic interaction through the global conductance *g*(*t*).

In (3) above, Δ is a measure of the variation in the natural frequency of the individual neurons which are described collectively by the NMM; specifically Δ is the width of a Cauchy distribution for the natural frequencies. The population excitability is *η*_0_, the centre of the Cauchy distribution. *A*(*t*) models the external drive and is a filtered version of the input *s*(*t*), defined by

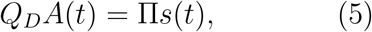

where *s*(*t*), the external stimulus, is a linear BSpline interpolation of the envelope, Π is the stimulus amplitude scale and

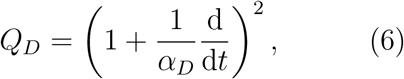

where *α*_*D*_ is a timescale for the filtering.

The synaptic input is also defined using a filter:

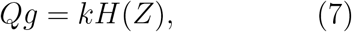

where

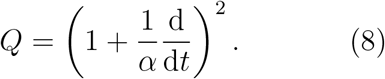

Here, *α* is another smoothing timescale, *H*(*Z*) is the measure of neuronal activity,

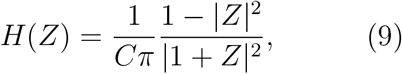

and *k* is the coupling strength inside the neural mass. For a full derivation, see Byrne et al. (2017).

We used two parameter sets for the NMM, which we refer to as the slow and fast models. For the slow model we increased *C*, and lowered both Δ and *η*_0_ relative to the parameters in the original paper Byrne et al. (2017). The values were chosen so that the model does not oscillate when there is no stimulation but oscillates at approximately 4Hz under slight stimulation. The fast model is identical to the model used by Byrne et al. (2017), producing approximately 15Hz oscillations. Using this original parameter set allows us to investigate what happens if the NMM’s base frequency is distinct from and higher than the stimulus frequency (typically 4Hz). Table 1 provides the two parameter sets.

**Table 1:** Parameters for the Slow and Fast NMM models.

| Parameter | $C$ F | $\Delta$ mA | $\eta_0$ mA | $v_s$ mV | $k$ | $\alpha$ ms<sup>-1</sup> | $\alpha_D$ ms<sup>-1</sup> |
| --- | --- | --- | --- | --- | --- | --- | --- |
| Slow ≈ 4Hz | 0.066 | 0.25 | 1.5 | -10 | 0.105 | 1/0.035 | 1/0.0056 |
| Fast ≈ 15Hz | 0.033 | 0.5 | 21.5 | -10 | 0.105 | 1/0.035 | 1/0.0056 |

The stimulus amplitude scale Π for the NMM, used to investigate the effect of varying drive strength, was set with reference to the excitability parameter *η*_0_, Π = *η*_0_*D*, where *D* is the drive-excitability-ratio. This allowed us to consider a range of drive strengths that were scaled to the particular internal excitability of the slow and fast models. The mean firing rate generated by the NMM was used to represent its activity. This firing rate is obtained through a conformal transformation of the complex Kuramoto order parameter *Z* to a new order parameter *W* = *πCr* + *iV*, where *r* is the firing rate, *V* is the mean membrane potential (Montbrió et al., 2015), and

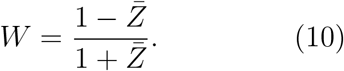

The firing rate is therefore

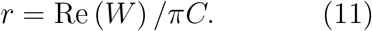

Initial conditions for *Z* ∈ ℂ were created by sampling its argument arg(*Z*) ~ Uniform (− *π, π*) and magnitude |*Z*| ~ Uniform (0.0, 0.9) such that it remained in the unit disk centred on the origin. Initial values for *g*, 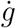, *A*, and 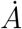were 0.

### Phase-resetting oscillator model

Our second model is the two-dimensional oscillator used by Oganian et al. (2023) to model phase-resetting. This contrasts with the biophysically derived NMM by representing the algorithmic process of phase-resetting, rather than the biological circuitry that may implement it. It is an oscillator with a fixed natural frequency that reacts to a stimulus only by shifting its phase towards a particular setpoint. The model consists of a Hopf normal-form with external stimulus *s*(*t*) that is modulated by the oscillator’s phase,

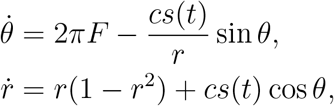

with phase *θ*, magnitude *r*, and coupling strength *c*.

This model orbits around the unit circle, with the stimulus forcing a phase-reset towards *θ* = 0. For this model the 10s stimuli streams were normalised to avoid the effect of varying power in the stimulus. Specifically, for the 60 trials corresponding to a given condition and test, the amplitude was scaled to have a mean per-stream-integral of 20000 over the samples in the interval [5.0, 10.0]s of the streams. This was simply a value close to the original integral of the first vowel condition trial. In principle, it would be possible to build a multi-oscillator version of this model, however, as for the NMM, a version with only a single macro-scale oscillator was tested. We set *c* such that the maximum possible phase correction was equal to *π*; this maximum occurs when *θ* = ±*π/*2 at the maximum value of *s*(*t*), assuming *r* = 1. Thus *c* varies according to the stimulus condition in order to maintain a consistent total “power” of the stimulus across tests.

The activity of the phase-resetting oscillator was taken to be *r* cos *θ* following the approach of Oganian et al. (2023). Initial conditions were sampled from random locations on the unit circle, *θ* ~Uniform (−*π, π*) and *r* = 1.

### Evoked response model

Our final model was a model of evoked activity. Unlike the NMM or phase-resetting oscillator, this model contains no self sustaining dynamics, instead it acts as a passive filter of an incoming stimulus. The model captures the direct effect of a stimulus on synaptic currents but, unlike the NMM, it does not include any representation of internal circuit dynamics. To ensure consistency between the stimulus processing of the NMM and the evoked model, we use the same linear differential operator *Q*, Equation (8) corresponding to the *α*-function response incorporated in the NMM by Byrne et al. (2017) as a synaptic filter. As such, the response of this model *g*(*t*) to stimulus *s*(*t*) is the solution of

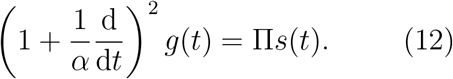

For an impulse-like stimulus, the model responds with a single alpha-function curve with rise time 1*/α*, thus providing a stereotypical evoked response. For a continuous stimulus, this is equivalent to a convolution with an alpha function kernel. We set *α* such that the rise time was 30ms. The drive strength Π was set to 1, without loss of generality, as the model is linear with respect to *s*(*t*). Scaling using Π therefore simply rescales the output without affecting its temporal structure. Since we are using the phase-based inter trial phase coherence (ITPC) for our evaluation, varying Π does not alter the results. Initial conditions were set such that *g* ~ Uniform (0, 1) and 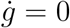.

### Sharpness Specific Tuning of Tracking

In Cucu et al. (2022), it was demonstrated that the EEG response to syllables is correlated with phoneme sharpness (see, e.g., fig. 6 in their paper). Here we use the same stimuli to investigate whether this experimentally-observed correlation can be recreated *in-silico*, fig. 1. The following first describes the experimental stimuli, then details how we constructed our in-silico stimuli. We then describe the experiments conducted using these stimuli. Finally, we detail parameter sweeps of noise level and drive strength.

**Figure 1:**
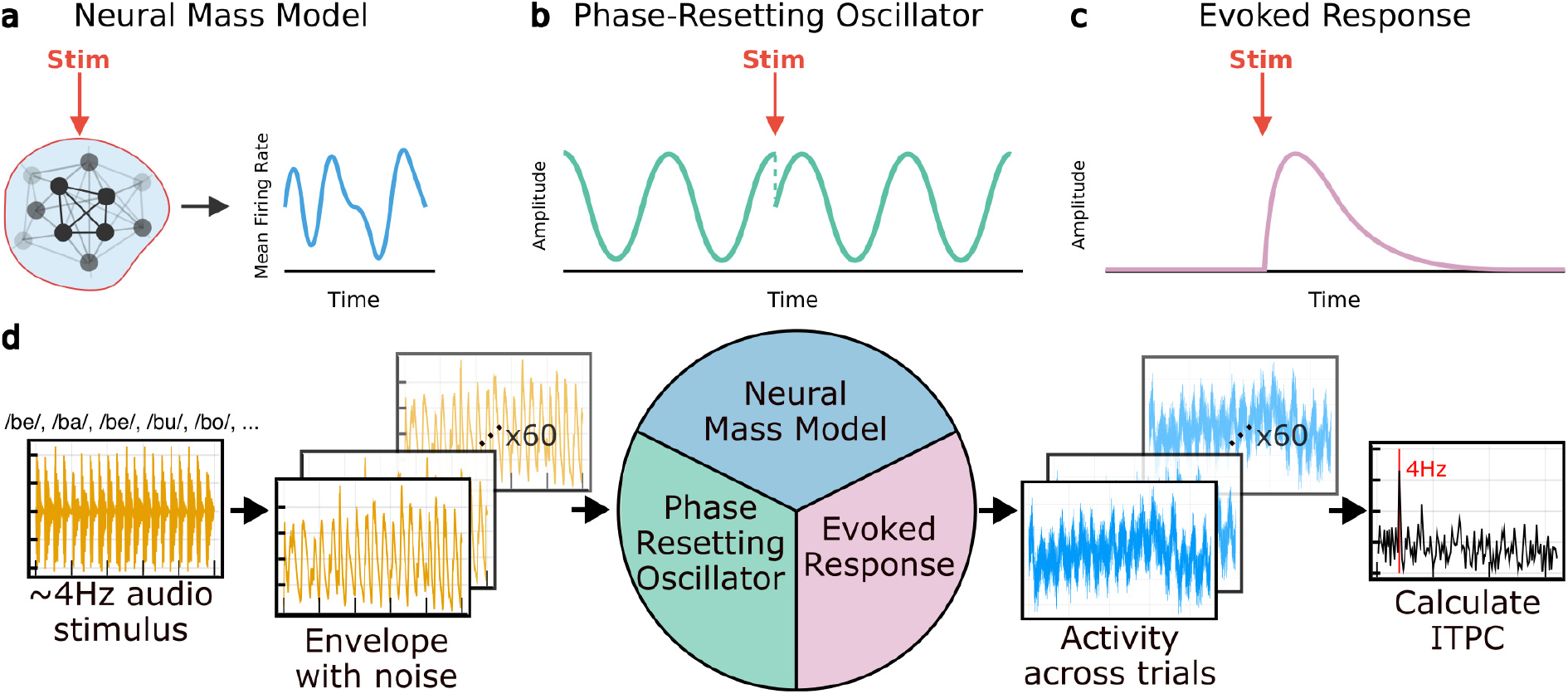
The three models used and the primary simulation design. **a**: a biophysical neural mass model. This is an exact mean field reduction of a network of spiking neurons, with state corresponding to the mean firing rate of the population. A stimulus is applied to the population as a whole. **b**: a phase-resetting oscillator. This is a phenomenological benchmark model of one of the hypothesised mechanisms of neural speech tracking. Stimuli act to reset the phase of a state that oscillates at a fixed natural frequency. **c**: an evoked-response model representing the alternative hypothesised mechanism of speech tracking. The model is silent until a stimulus is applied. For an impulse stimulus, a stereotypically shaped response is produced. **d**: schematic of our reproduction of Cucu et al. (2022)’s EEG experiment *in-silico*. Raw audio stimuli, consisting of near-isochronous 4Hz consonant-vowel syllable sequences, are processed into envelopes with noise added to simulate trials. These are used to drive the three models. The inter-trial phase coherence is computed for each model across the trials. The 4Hz ITPC is then used to investigate the effect of phoneme sharpness on tracking strength. Example model activities are presented in fig. 1-1 in the exended data.

#### Experimental Stimuli

The stimuli are near-isochronous 4Hz auditory streams of consonant-vowel (CV) pairs from 15 different conditions (Cucu, 2021). The 15 conditions are 14 consonant conditions, where the consonants are paired with random vowels in the streams, as well as a vowels only condition. For each condition, there are three distinct auditory streams corresponding to different random vowel sequences that the consonant is paired with, these are hereafter referred to as ‘items’. For example, one of the ‘k’ items begins

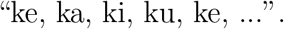

As such, there are 45 items in total (15 conditions, three items per condition).

#### Stimulus construction

The experimental audio stimuli were transformed to form sound envelopes for each stream. Following the processed used in Cucu et al. (2022) to compute the sharpness of the phonemes, we passed the raw auditory data through a cochlear band-pass filter, with frequencies ranging from 80 to 8000Hz in 32 log-spaced bands and computed the Hilbert transform to get the analytical signal in each frequency band. These were then summed to give the final envelope. The maximum amplitude of the envelopes was approximately 0.25.

Additive syllable-based noise was included to simulate background “chatter” during the experiment. The noise streams were constructed by squeezing and stretching the existing syllable streams. For each instance of the noise, 20 syllables were selected at random, and these were concatenated and subject to temporal distortion while keeping the overall stimulus length fixed. We produced 60 different instances of this noise, giving 60 trials for each item. To ensure transient effects from the drive were minimised we appended five seconds of syllable-based noise of the same sort to the beginning of each stimulus stream, aligned such that the end of a distorted syllable aligned with the start of the stimulus stream. Overall there were thus 60 trials ×3 items for each of 15 conditions giving, in total, 2700 distinct 10s in-silico stimuli. We explored the effect of the strength of the noise using a noise-to-stimulus ratio *λ*. For a particular noise-to-stimulus ratio *λ*, the stimulus envelopes were multiplied by (1 − *λ*), before the noise scaled by *λ*, was added to them. This therefore required 2700 stimuli streams per *λ* setting.

For the evoked model we investigated the response to three variants of the auditory stimuli. These variants were: the original envelopes as used for the NMM and phase resetting model, the derivative of a 10Hz low-pass filtered version of the envelopes, and a series of impulses corresponding to peak-rate events. 10Hz was selected for the low-pass filter as it is the upper end of where the power in the speech modulation spectrum is concentrated (Ding et al., 2017b). We used peak-rate impulses because peak-rate acoustic edge representations have been identified in auditory speech cortex Oganian and Chang (2019).

To create the peak-rate impulse stimuli, we smoothed the original envelopes with a 10Hz low-pass filter and then applied a peak finding algorithm on the derivative of the envelope. This identified the peaks, with a lower threshold of 0.1 peak magnitude. Once the peaks were identified, impulses were created at the time points of the peak events, with heights set to the magnitude of the derivative at that point. The impulses were then scaled to a range of [0.5, 1.0] following the approach of Oganian et al. (2023).

#### Experiments

Using the trial stimuli outlined above, we conducted several in-silico experiments to investigate the response of the models and simulate Cucu et al. (2022)’s experiment.

Specifically, we applied the stimuli to the models as an external drive, and recorded the response. The ITPC was calculated over the 60 trials for a given condition item. For each condition, the mean of its three items ITPCs was then calculated. To compare to the experimental results in Cucu et al. (2022) we then calculate the correlation between this mean ITPC and the phoneme sharpness.

We used latency to maximum derivative, *L*, to measure phoneme sharpness as this was identified to be most strongly correlated with tracking strength by Cucu et al. (2022) amongst a range of acoustic edge features. We used the values derived by Cucu et al, as shown in Table 2. These were obtained by identifying the maximum derivative of the envelope after applying a band pass filter between 2 and 10Hz, and then identifying the time taken to reach this point from the onset of the phoneme.

**Table 2:** Latency to peak derivative, *L*, for each stimulus condition. Cells are ordered by increasing *L* and coloured according to the phoneme type: vowel, voiced stops, unvoiced stops, nasals, sibilants, liquids, and fricatives.

| Condition | b | n | m | d | vowel | l | g | p | t | k | r | v | z | f | s |
| --- | --- | --- | --- | --- | --- | --- | --- | --- | --- | --- | --- | --- | --- | --- | --- |
| **L (ms)** | 17.01 | 18.59 | 19.19 | 21.63 | 23.49 | 29.98 | 29.17 | 37.01 | 47.66 | 51.07 | 52.77 | 64.88 | 87.62 | 93.07 | 98.83 |

All models were tested with the envelope stimuli. The evoked model was also tested with the derivative of the envelope, and peak-rate impulse stimuli. We ran these tests for a range of noise-to-stimulus ratios and, for the NMM, we also investigated the effect of the drive strength in relation to the excitability of the model (See the “Simulation settings: noise and drive strength” section below). An example response of each model is presented in the extended data, fig. 1-1.

#### Simulation settings: noise and drive strength

To identify reasonable simulated experimental conditions for our tests, in particular the noise level and drive strength, we calculated the concordance correlation coefficient between the model ITPCs and those obtained in Cucu et al. (2022)’s experiment. We wanted to find operating conditions that produced tracking of a similar strength to that observed in Cucu et al. (2022), as well as a similar correlation with phoneme sharpness, taking this to mean we were operating under ‘realistic’ experimental conditions. To be clear, this is not to say we were attempting to fit to biologically plausible values of these parameters, only to approach dynamical regimes that appear qualitatively similar to the response in experiment. The *D* and *λ* parameters effectively represent input signal gain, and background noise levels, and can be thought of as tuning the experimental environment, rather than tuning the behaviour of the models themselves.

Specifically, we tested all of the models using noise-to-stimulus ratios *λ* ∈ [0, 0.05, 0.1, 0.3, 0.5, 0.7, 0.9, 1.0]. This ratio was used in the stimulus construction to determine how much noise to add to the stimulus stream. For instance, with *λ* = 0.6 the drive is 60% noise and 40% stimulus. For the NMM we also varied the drive strength. We did this relative to the excitability, *η*_0_, of each of the fast and slow models. To do this, we produced a drive-excitability-ratio parameter *D*, taking log-spaced values between 0.1 and 300, but capped at 100 for numerical stability. As such *D* ∈ [0.1, 0.496, 2.46, 12.2, 60.5, 100.0]. Using this, we then set the drive scale parameter Π = *η*_0_*D*, where *η*_0_ was the excitability of the particular fast or slow model being tested; see the NMM model description, Equation 5, for further detail. For the phase-reset and evoked-response models, the drive was not varied. For the phase-reset model we instead used normalisation of the stimuli and model parameters to set a consistent scale of response, meaning any changes to the drive scale would get normalised out. For the evoked model the drive strength was fixed to 1, as varying this does not affect the purely phase-based ITPC evaluation. To be clear, the *D* and *λ* parameters effectively represent input signal gain, and background noise levels, and can be thought of as tuning the experimental environment, rather than tuning the behaviour of the models themselves.

#### Post-hoc noise

In addition to the noise in the stimulus, constructed out of distorted syllables, we added post-hoc 1*/f* noise to all of the model activities with a signal to noise ratio of 0.5. This was used to simulate the noise present in EEG. We used the PowerLawNoise.jl (v0.0.1) package to produce the noise (https://github.com/astrogewgaw/PowerLawNoise.jl).

### Model Dynamics Investigation

Once we had established that the NMM, as well as the phase-resetting oscillator and evoked model, could reproduce the result from Cucu et al. (2022), we ran further experiments to explore the models’ dynamics more closely. First, we used the PCM, which was developed to distinguish between evoked or phase-reset-like responses to rhythmic tones. Then, we re-ran the CV-syllables experiment while varying the syllabic rate of the stimuli. This is a similar test, but reports changes in tracking strength only, rather than the changes in phase-lag between stimulus and response that the PCM reports. Finally, we calculated the IEPC of the NMM’s response over a time window centred about the phoneme onsets of a particular condition stream to identify any transient, onset-related dynamics.

#### Phase Concentration Metric Experiment

The PCM evaluates the similarity of phase-lags between the stimulus envelope and model response across stimulus rates. The phase lag is computed from Gaussian filtered versions of the stimulus and response, see the *Evaluation metrics and statistical analyses* section for details. The PCM varies between zero and one as it represents the magnitude of the circular mean of unit vectors in the complex plane. For an evoked-response model, the PCM is expected to be low, as the phase lag shifts with the stimulus rate. For a phase-resetting oscillator, the PCM is expected to be high as the phase lag remains more consistent under stimulus rate changes.

To use the PCM to investigate the behaviour of the NMM, we generated sine wave stimuli of varying frequencies and applied a DC shift of +1 to ensure they were strictly positive. These were used to drive the NMM and phase-resetting oscillator. For the evoked model, unit impulses timed at the peaks of the sine wave were used instead. We varied the stimulus frequency from 2Hz to 15Hz in 60 steps and computed the PCM from the resulting trajectories. Details of the PCM implementation are provided in the Evaluation Metric and Statistical Analysis section below. Model parameters were kept the same as for the sharpness-tuning experiment, except the drive amplitude for the NMM, which was fixed at 15.0.

#### Speech Rate Experiment

To vary the syllabic rate of our stimuli, we scaled the time used to sample from the stimulus interpolation object storing them. In the ODE simulations, the drive was obtained from an interpolation object that returned a value for a given time, i.e. *drive* = stimulus interpolator (*t*). When sampling using the actual time, this provided the 4Hz stimulus, which was 10s long. Here, we instead sampled using the real time *t* scaled by a factor *τ*. For instance, for a run with a desired syllabic rate of 1Hz, *τ* = 0.25, and for a syllabic rate of 12Hz, *τ* = 3. This necessarily results in different length stimuli, for instance, the 10s baseline duration would become 5s when sampling at twice the rate for a 8Hz syllabic rate. To ensure that the calculation of ITPC was done over a suitable number of cycles, we extended the stimulus period from [5, 10]s to [5, 20]s, simply repeating the compressed or stretched stimulus segment as many times as necessary. This repetition was also done separately for the 5s of noise that prepended the stimulus to ensure that the noise did not enter the ITPC calculation window. Thus, to avoid transient effects, we set the ITPC calculation window to [5.5, 20]s, skipping the first 0.5s of the stimulus period.

As we were interested only in how the syllabic rate affected the tracking strength, we kept other parameters constant. For the simulation settings, we chose *D* = 12.2 as this was found to be optimal for the sharpness-tuning experiment and *λ* = 0.3 to ensure strong enough tracking to observe any variation due to syllabic rate. Model parameters were the same as those used in the sharpness-tuning experiments. We ran this test for 20 trials, for one item, of each of the conditions, reporting the mean ITPC across these 15 conditions for each syllable rate.

#### Inter Event Phase Coherence About Phoneme Onsets

Following the approach in Oganian et al. (2023), we computed the IEPC over a time window centred at the onset of the phonemes. Specifically, for one of the ‘b’ phoneme condition streams, we identified the peaks in the derivative of a 10Hz low-pass filtered version of the envelope that occurred between 5s (the start of the stimulus period) and 9s, taking these to be the phoneme onsets. This was done for 20 trials, in total, identifying 340 distinct phoneme onsets.

We band-pass filtered the response of the fast NMM to these 20 ‘b’ condition trials with a second order Butterworth filter into 100 frequency bands centred at frequencies uniformly spaced between 0.67 and 35Hz. Each *f* Hz band was set to *f* ± 1Hz, except for frequencies below 1Hz, where the lower bound was set to 0.1Hz. Zero-phase filtering was used to prevent phase distortion. We applied a Hilbert transform to the band-pass filtered signals to obtain the analytical signal. The instantaneous phase in each frequency band was then taken to be the argument of the analytical signal.

These instantaneous phase values were segmented into time windows centred at each of the 340 phoneme onsets. Each window ranged from −0.25 to +0.25s about the phoneme onset. Finally, we converted the instantaneous phase values into exponential form, creating complex unit vectors. The magnitude of the mean of these vectors gave the IEPC for each frequency band (see the *Evaluation metrics and statistical analyses* section for formal details). To further identify any transient effects of the phoneme onset, we computed a baseline adjusted IEPC. To do this, we subtracted the mean IEPC value between −0.2 to −0.1s in each frequency band from the original IEPC values.

Additionally, to explore whether any ‘ringing’ entrainment was present at the syllabic rate after stimulus cessation, we computed the 4Hz IEPC over all 60 of the ‘b’ condition trajectories from 0s to 20s, with the first 5s being noise, 5 to 10s the ‘b’ syllable condition stimuli, and with the drive amplitude set to 0 from 10s onwards.

### Evaluation metrics and statistical analyses

#### Inter Trial Phase Coherence

The ITPC of the model activity across the 60 trials for a given stimulus stream was used to measure the alignment to the periodic stimuli, as in Cucu et al. (2022); Ding et al. (2017a). What is called the ITPC here is often called the mean square resultant, *R*^2^, in more general context. In particular, for a given frequency *f* :

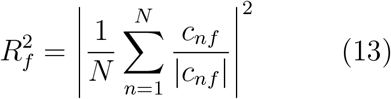

where *c*_*nf*_ is the *f* Hz Fourier coefficient of the *n*th signal, so that

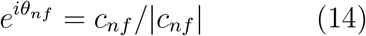

defines the phase *θ*_*nf*_. Here, *n* is the trial index with *N* trials in total. The mean of the ITPC at 4Hz across each condition’s test streams was calculated, and used in the computation of the correlation with the phoneme sharpness. The ITPC ranges between 0 and 1, where 0 indicates no coherence or alignment between the *f* Hz frequency components in the trial signals, and an ITPC of 1 indicates perfect coherence or aligned phase.

#### Correlation and Multiple Comparison Corrections

As in the original analysis, Cucu et al. (2022), we measure the correlation between the tracking strength, 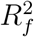 and the phoneme sharpness. We evaluate the significance of this correlation for each model using a two tailed *t*-test with 13 degrees of freedom (*n* − 2, *n* = 15 conditions). The effect of multiple comparisons is assessed and reported separately for both the Bonferroni correction and the Benjamini-Hochberg FDR correction (with a false discovery rate of 0.05). Confidence intervals are computed for the correlation coefficients using the z-transformation. We do this at the simulation conditions that we decided are most realistic through comparison of the mean ITPC across conditions of the models to that of the experimental EEG result (See “Simulation settings” section). However, in SI we report how the ITPCs, correlations, and confidence intervals change as we change the operating parameters for the simulation. Overall, we are seeking to make a qualitative comparison between the response of these models and the experimental EEG result.

#### Concordance Correlation Coefficient

To identify reasonable experimental conditions for our tests, for instance, noise levels and drive strengths, we compared the model ITPCs across conditions with those identified in experiment using the concordance correlation coefficient *ρ*_*c*_ (Lin, 1989),

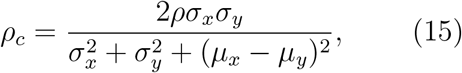

where *ρ* is the correlation coefficient, 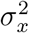 and 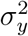 are the variances of the datasets, and *µ*_*x*_ and *µ*_*y*_ are the means of the datasets. *ρ*_*c*_ assesses the agreement between datasets, accounting for bias and scale as well as correlation by quantifying how far the data deviate from each other, or equivalently the line *y* = *x*, on a plot of *x* versus *y*.

#### Phase Concentration Metric

To compute the phase lag between the stimulus and response at a particular stimulus-rate *f*_*s*_, the time series were first filtered using a Gaussian kernel in the frequency domain with centre *µ* = *f*_*s*_ and width *σ* = *f*_*s*_*/*2.

If *X*(*f*) is the Fourier transform of the signal (the response, *r*(*t*) or stimulus envelope, *s*(*t*)), then the filter kernel,

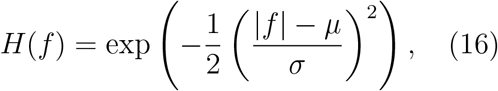

and the filtered signal *x*(*t*) is obtained via the inverse Fourier transform of the point-wise product:

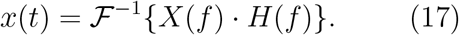

The instantaneous phases, *ϕ*_*r*_(*t*) and *ϕ*_*s*_(*t*), are then calculated by applying a Hilbert transform to the filtered signals. The phase lag time series then follows as

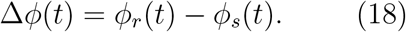

Finally, these phase lags are transformed into complex unit vectors and their mean taken to produce a mean phase lag vector for the stimulus rate,

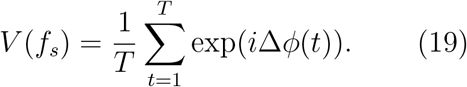

The PCM is then the magnitude of the mean of these vectors across stimulus rates:

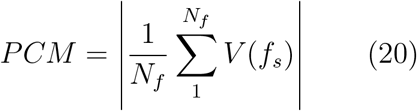

#### Inter Event Phase Coherence

The fundamental calculation of the IEPC is similar to that used for the ITPC, however there are several key differences. For the ITPC, we calculate the circular mean of complex unit vectors defined by the *f* Hz Fourier coefficients across *N* trials for a given stimulus stream. For the IEPC, we instead calculate the circular mean of complex unit vectors obtained from the instantaneous phase of Hilbert transformed band-pass filtered signals, across the *N* = 340 time windows centred at the phoneme onsets. Thus for a set of instantaneous phases *ϕ*_*nf*_ (*t*) from the analytic signal in frequency band *f* over windows *n*∈ [1, *N*] we have

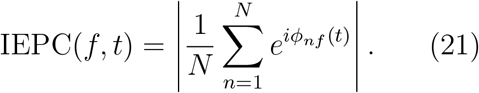

### Code Accessibility

The code/software and data described in the paper is freely available online at https://github.com/Modelling-Speech-Processing/Shannon_et_al**redacted**, and the code is also provided in Extended Data 1.

## Results

At representative operating conditions (*λ* = 0.9, *D* = 12.2), both the NMM and the phase-resetting oscillator recapitulate the correlation from Cucu et al. (2022), (*r*(13) = −0.91, *p* < 0.001), fig. 2**a**-**c**. This is the case for both the slow, (*r*(13) = −0.85, *p* < 0.001, 95% CI [−0.95, −0.61]) and fast (*r*(13) = − 0.91, *p* < 0.001, 95% CI [−0.97, −0.74]) parameter combinations of the NMM. Overall, the ITPC is greater for the slow version of the model. This may be due to the alignment of the baseline oscillations with the frequency of the 4Hz stimulus. The phase-resetting oscillator model also successfully reproduces the negative correlation when driven by the envelope: *r*(13) = −0.79, *p* = 0.0005, 95% CI [−0.93, −0.46].

**Figure 2:**
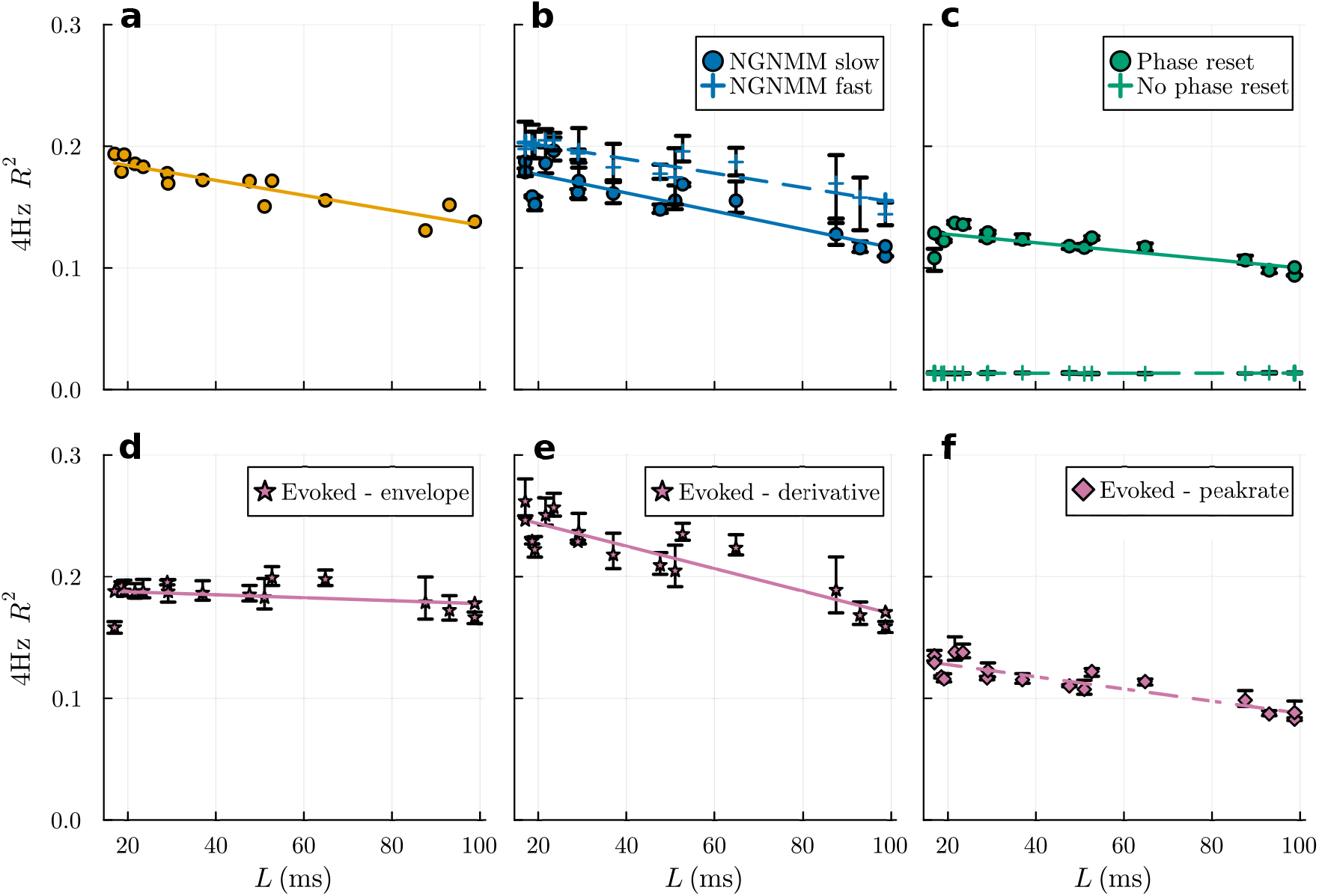
Mean 4Hz ITPC across test streams vs envelope sharpness as measured by the latency to maximum derivative *L*. Each point represents a different consonant-vowel condition. **a**: experimental EEG result from Cucu et al. (2022). **b**: 4Hz coherence of the NMM response under the two parameter settings, fast (≈ 15Hz) and slow (≈ 4Hz). **c**: 4Hz coherence of the phase-resetting oscillator response, with and without phase-resetting. **d**: 4Hz coherence of the evoked response to the stimulus envelope. **e**: 4Hz coherence of the evoked response to the derivative of the envelope. **f** : 4Hz coherence of the evoked response to the peak-rate event impulse train stimuli. Error bars show the range of the 4Hz ITPC over the three test items for each condition.

The evoked model does not respond in the same way to the stimulus-envelope. Unlike for the NMM and phase-reset models, it does not create a negative correlation between sharpness and tracking strength in the response of the evoked model, *r*(13) = −0.30, *p* = 0.270, 95% CI [−0.71, 0.25], (fig. 2**d**). This is despite still causing the 4Hz activity to align across trials, with a mean 4Hz ITPC= 0.184. However, the evoked model does produce the negative correlation observed experimentally when driven by the pre-processed forms of the stimulus. Specifically, with the derivative of the stimulus envelope, *r*(13) = −0.88, *p* < 0.0001, 95% CI [−0.96, −0.67] and with the peak-rate impulse stimulus, *r*(13) = −0.87, *p* < 0.0001, 95% CI [−0.96, −0.64], fig. 2**e** and **f**. The evoked model demonstrates the negative correlation for all but the envelope-based stimulus. Overall, under this set of simulation conditions, *λ* = 0.9, and, for the NMM, *D* = 12.2, the magnitude of the 4Hz ITPCs closely resemble the experimental result.

Note that the high amount of noise in the stimulus was required to obtain this similarity. This is made evident by observing the concordance correlation coefficient between the EEG based ITPCs and each model’s ITPCs as the simulated noise strength *λ* is increased, fig. 3**c**.

**Figure 3:**
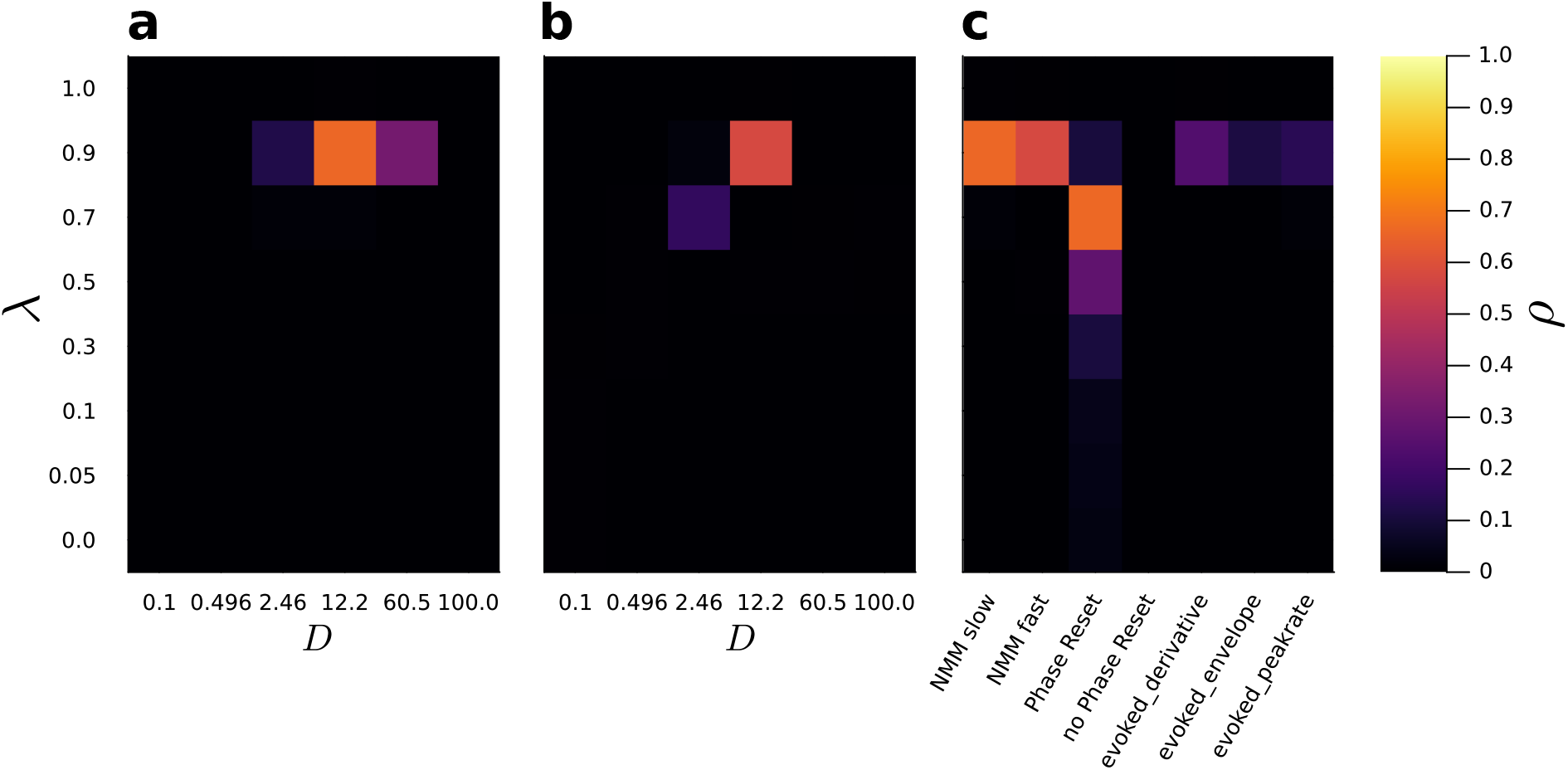
Heatmaps of the concordance correlation coefficient, *ρ*, between the mean ITPC, *R*^2^, per phoneme condition of the models and the EEG data. **a, b**: the slow and fast NMM respectively, as *D* and *λ* are varied. **c**: for each model as *λ* is varied. The admissible range of *ρ* values is between 0.0 and 1.0, with 1.0 indicating a perfect match between the models *R*^2^ values and those obtained from the EEG (Cucu et al., 2022). The full, per-phoneme-condition, ITPCs for each model under every experimental condition are presented in fig. 3-1 and fig. 3-2, alongside the corresponding correlations and *p* values in tables 3-1 to 3-4.

For the NMM and evoked model, *ρ* is highest when *λ* = 0.9, reaching 0.66, 0.57, and 0.24 for the slow NMM, fast NMM, and derivative driven evoked model respectively. For the phase resetting oscillator *ρ* is highest at *λ* = 0.7, reaching 0.67. Of course, for *λ* = 1.0, 100% noise, the tracking strength becomes negligible, and any correlation with phoneme sharpness is obscured. For lower values of *λ* we found that the ITPCs became too high. At a point that varied between models (fig. 3-2), the 4Hz ITPCs saturated close to 1.0, indicating complete coherence, and again obscuring any correlation with phoneme sharpness. This is the cause of the very low values of *ρ* for lower noise levels. The phase resetting oscillator required slightly more signal strength to entrain, resulting in the peak *ρ* at *λ* = 0.7 and the lower 4Hz ITPC’s in fig. 2 at *λ* = 0.9 in comparison to the experiment. The correlation is still present however. The other experimental condition we needed to set was the drive strength for the NMM. Figure 3**a** and **b** present the concordance correlation coefficient between the EEG based ITPCs and the slow and fast NMM’s ITPCs respectively as both the drive strength, *D*, and noise level, *λ*, were varied. It is clear that *D* = 12.2 was the optimum drive-excitability ratio for accurate reproduction of Cucu et al’s correlation. The full set of ITPCs for each phoneme condition for the slow and fast NMM models across *D* settings are presented in the extended data, fig. 3-3.

All of the models except the non-phase resetting oscillator, and the envelope-driven evoked model produce significant uncorrected correlations (*p* < 0.001, Table 3). This remains the case when corrected using a Benjamini-Hochberg FDR correction with a false discovery rate of 0.05. Under the more stringent Bonferroni correction, the phase-resetting oscillator correlation no longer reaches *p* < 0.001. A full set of correlations and related statistics for all models across all *λ* and *D* settings are provided in the extended data, tables 3-3 to 3-4.

**Table 3:** Correlation between phoneme sharpness and 4Hz ITPC for each model variant and the original data obtained by Cucu et al. (2022). We present uncorrected and corrected *p* values for the correlations. Bonferroni and Benjamini-Hochberg FDR corrections were calculated for eight comparisons (the model and stimulus variants) and a false discovery rate of 0.05.

| Model | Version | $r$ | 95% CI | $p$ | $p_{\text{Bonferroni}}$ | $p_{\text{FDR}}$ |
| --- | --- | --- | --- | --- | --- | --- |
| EEG | – | −0.913 | −0.97, −0.75 | $2.0 \times 10^{-6}$ | – | – |
| NMM | Fast | −0.909 | −0.97, −0.74 | $2.7 \times 10^{-6}$ | $2.2 \times 10^{-5}$ | $2.2 \times 10^{-5}$ |
| | Slow | −0.854 | −0.95, −0.61 | $5.2 \times 10^{-5}$ | $4.1 \times 10^{-4}$ | $1.0 \times 10^{-4}$ |
| Oscillator | Reset | −0.786 | −0.93, −0.46 | $5.2 \times 10^{-4}$ | 0.0042 | $6.9 \times 10^{-4}$ |
|  | No reset | 0.165 | −0.38, 0.62 | 0.56 | 1.0 | 0.56 |
| Evoked | Envelope | −0.303 | −0.71, 0.25 | 0.27 | 1.0 | 0.31 |
| | Derivative | −0.879 | −0.96, −0.67 | $1.6 \times 10^{-5}$ | $1.3 \times 10^{-4}$ | $6.4 \times 10^{-5}$ |
| | Peakrate | −0.870 | −0.96, −0.64 | $2.5 \times 10^{-5}$ | $2.0 \times 10^{-4}$ | $6.8 \times 10^{-5}$ |

It is clear that the NMM produces ITPCs that most closely resemble the experimental ITPCs (fig. 4) for this set of simulation conditions (*λ*= 0.9, *D*= 12.2). However, although overall the model shows a roughly correct negative correlation between sharpness and tracking strength, on a per phoneme basis there is still some disagreement on the exact coherence, as well as a difference in the exact ordering of phonemes. For instance, the vowel condition has the third highest ITPC for the EEG experimental result, but creates the highest ITPC for the slow NMM. The phase-resetting model would more closely match the EEG result with *λ*= 0.7 (see fig. 3), however we are comparing the models here under the same noise condition. The main point of disagreement between the experimental and simulated NMM model results is in the ITPC of the ‘p’, ‘m’, ‘n’, ‘z’, and ‘v’ conditions. The successful recreation of the negative correlation between sharpness and tracking strength seems to be driven by the ITPC of the models being accurate, in their coherence relative to the other conditions, for the vowel and sibilant conditions that lie at either end of the sharpness scale.

**Figure 4:**
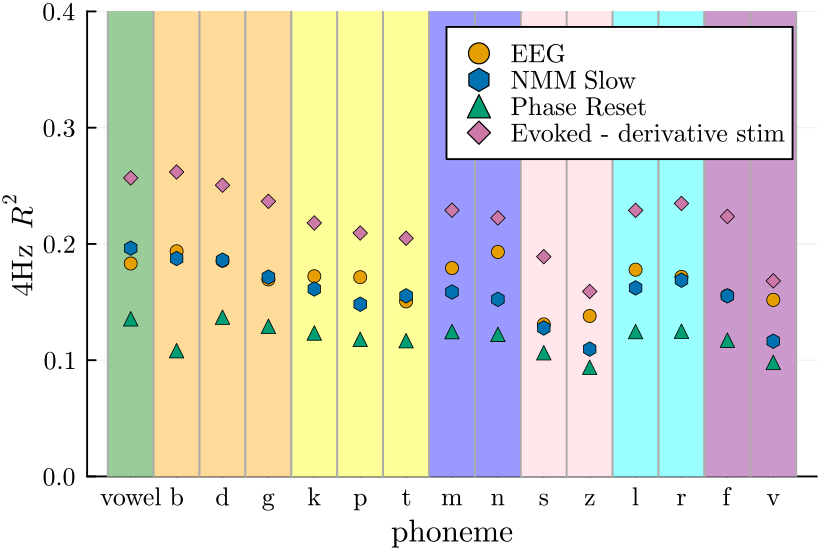
Mean 4Hz ITPC vs phoneme condition for each model alongside the experimental EEG result. Each point represents a different consonant-vowel condition. Vertical bands are coloured according to the phoneme type: vowel, voiced stops, unvoiced stops, nasals, sibilants, liquids, and fricatives.

Although all three models seem to be able to reproduce the correlation from Cucu et al. (2022), their mechanisms differ. The two phenomenological models have known mechanisms that are inherent to their construction, these being evoked responses and phase-resetting. However to understand the NMM’s behaviour further, we carried out investigations into what kind of mechanisms support its tracking of the CV-syllable stimuli.

The first of these tests was the PCM test, assessing how the phase lag between the stimulus and the model response’s stimulus-rate frequency component changes with the stimulus rate. The PCM was constructed such that an evoked model would exhibit a mean phase lag that changes as the stimulus frequency changes, and such that oscillating models that phase reset have a more consistent, concentrated phase lag as the stimulus frequency changes (Doelling et al., 2019). Our evoked model and phase-resetting oscillator demonstrate this (fig. 5**a** and **b** respectively). Using these as a baseline example of possible behaviour, we can see that both the slow and fast NMM’s look most similar to the evoked model. In particular, although all the models display a systematic relationship between phase-lag and frequency, this has higher variance for the NMM and evoked models.

**Figure 5:**
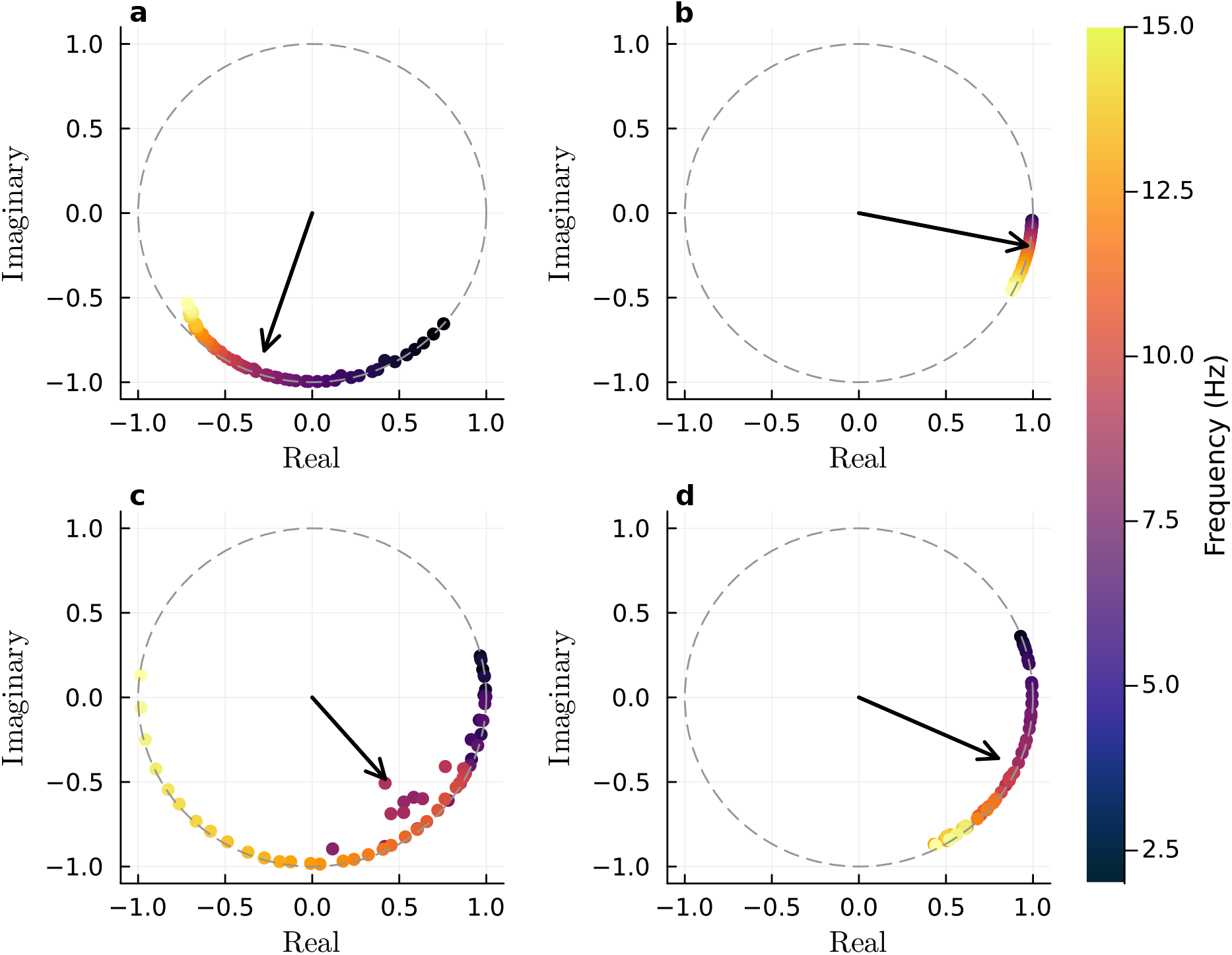
Phase Concentration Vectors for each model in the PCM test. **a** the evoked model. **b** a tuned phase-resetting oscillator. **c** the slow NMM. **d** the fast NMM. Each vector, represented by a dot, is the mean phase-lag across a single response to the stimulus, at a frequency defined by the colour. The phase-resetting oscillator, and NMM’s were driven by sine waves. The evoked model was driven by unit impulses aligned to the peaks of the same sine waves. Evoked responses result in a shifting phase lag as stimulus frequency changes. Phase-resetting results in a more consistent phase lag. The PCM, the magnitude of the circular mean of the phase concentration vectors, is indicated by the black arrows.

**Figure 6:**
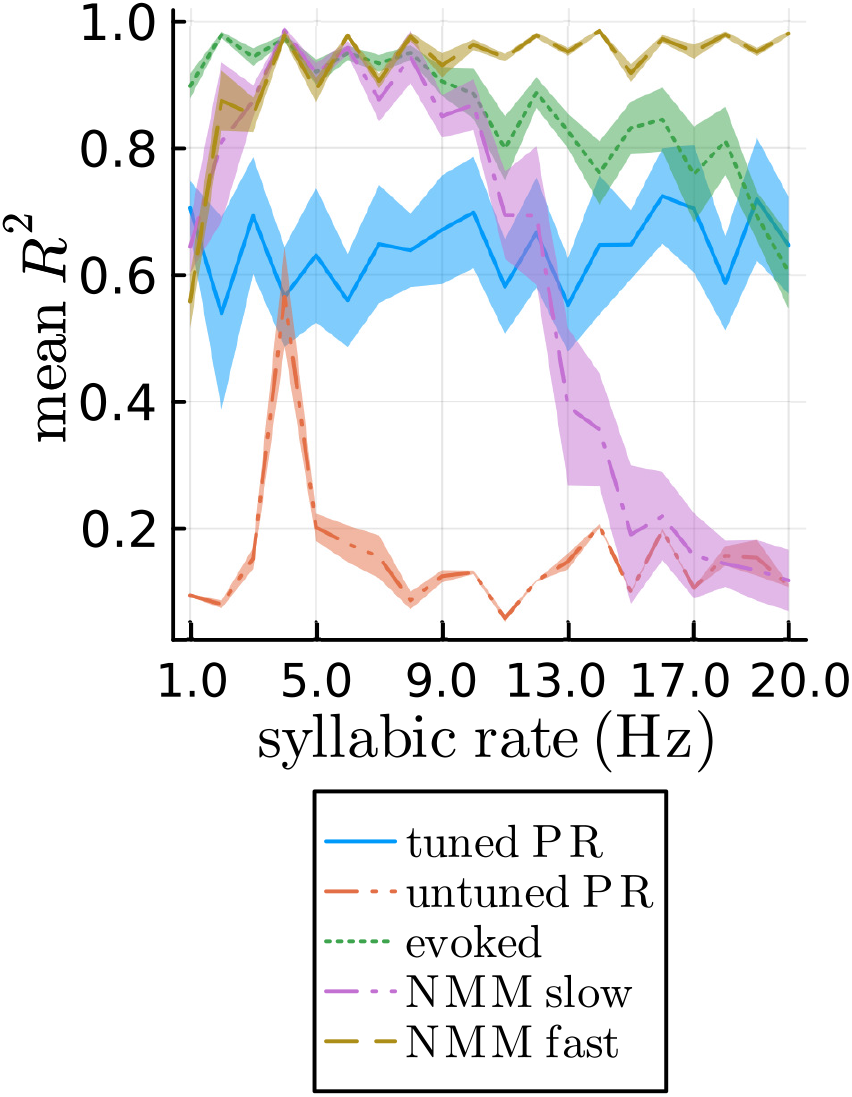
Mean ITPC, *R*^2^, across all phoneme condition items, as the syllabic rate is varied. Blue line, a phase-resetting oscillator tuned so that its own natural frequency matches the syllabic rate, Red line, a phase-resetting oscillator with 4Hz natural frequency. Green line, the evoked model. Purple line, the slow NMM. Gold line, the fast NMM. Shaded regions represent one standard deviation.

We also examined how the models respond to different syllable rates by using the CV-syllable based drive with a varying rate. Here we report the mean *R*^2^ at the stimulus rate across 60 trials across all conditions, similar to the sharpness specific tuning experiment. However we have manipulated the syllabic rate to values between 1 and 20Hz, fig. 6. The ‘untuned’ phase resetting oscillator, with a fixed natural frequency of 4Hz shows a clear peak in coherence at 4Hz as expected. When this model is set such that it oscillates at the syllabic rate, it successfully entrains across all rates, showing high *R*^2^ across all rates. In contrast, the evoked response model entrains strongly at low frequencies and maintains this across the tested frequencies with a slight fall as the rate increases. At higher frequencies, the low ITPC is possibly due to the timescale of the evoked response being too large for these frequencies. Interestingly and in contrast to the PCM experiment, the slow and fast NMM’s no longer resemble the evoked model. Instead, the slow NMM has a peak centred at its natural frequency like the untuned phase-resetting oscillator, but much broader, and the fast NMM demonstrates high ITPC across all syllabic rates. This is most similar to the tuned phase-resetting oscillator, despite the NMM not having its natural frequency tuned to match the syllabic rate.

The PCM and syllabic rate tests provide conflicting conclusions. The PCM test suggests that the slow and fast NMMs produce evoked responses. The syllabic rate test suggests that the slow NMM approximates an untuned phase-resetting oscillator while the fast NMM is most similar to the tuned phase-resetting oscillator. To further characterise their behaviour, we inspected the NMMs’ responses to individual CV-syllables. Figure 7 presents the response of the fast NMM to the first syllable in the stimulus period, after 5s of syllable-based noise are applied. 20 trial trajectories are plotted with the mean trajectory overlaid. It is clear that before the phoneme onset at time 0.0s, the trajectories are desynchronised due to the noise. After the phoneme onset their phase is reset and they become temporally aligned. This indicates that the fast NMM is being phase reset by the onset of the phoneme envelopes. It seems that the fast oscillations are being gated-by or nested-within the 4Hz syllabic rate oscillations. The phoneme onsets also cause phase resetting in the slow NMM model, fig. 8. However, no nested oscillations or cross-frequency coupling occurs as the syllabic rate matches the natural frequency of this model. We use the terms cross-frequency coupling and gating here in the nonlinear dynamical systems sense rather than to describe interactions between separate neural populations. Specifically, this refers to nonlinear resonances between frequencies within a single state of a nonlinear system: the slow 4Hz drive with its sharp phoneme onsets elicit bursts of high frequency activity. These are temporally gated by the envelope of the syllables as the decrease in drive along the tail of the envelope eventually suppresses the fast oscillations.

**Figure 7:**
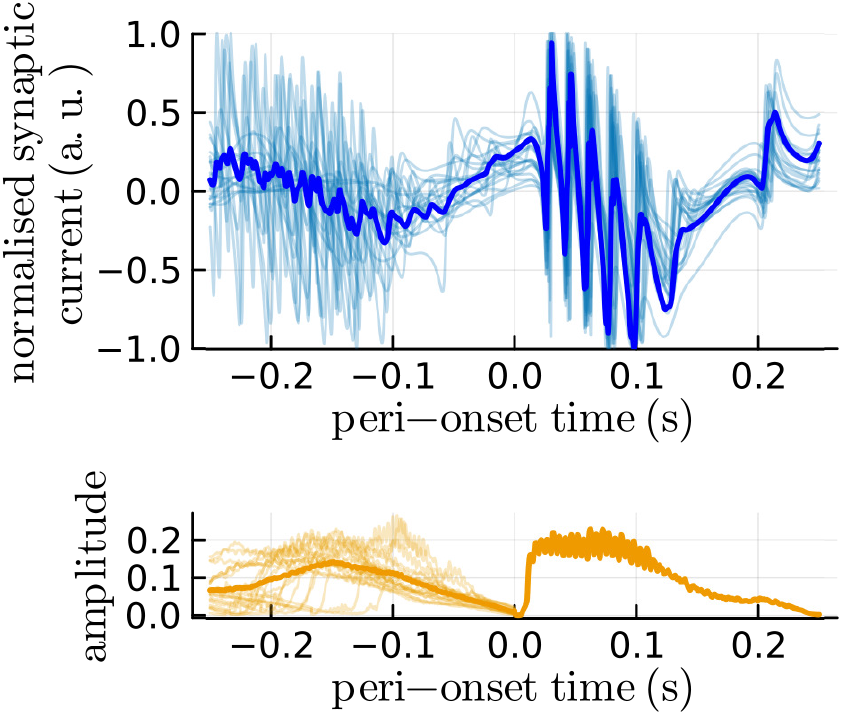
Response of the fast NMM to the initial phoneme onset over 20 trials of the ‘b’ phoneme condition. Despite having random phases at the onset of the stimulus, the trajectories align due to phase resetting. Noise-to-stimulus ratio *λ* = 0.0 for clarity.

**Figure 8:**
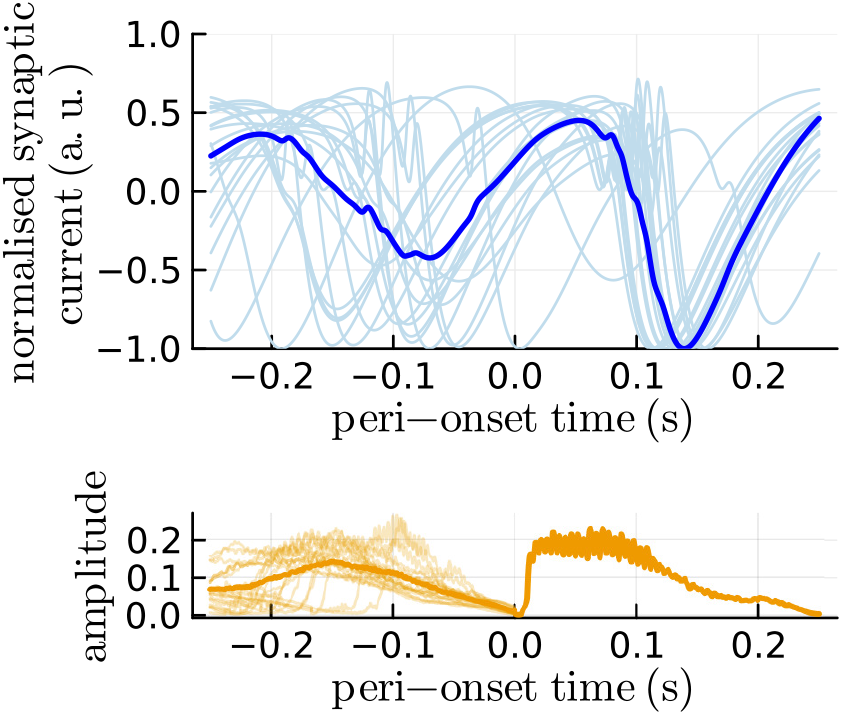
Response of the slow NMM to the initial phoneme onset over 20 trials of the ‘b’ phoneme condition. Despite having random phases at the onset of the stimulus, the trajectories align due to phase resetting. Noise-to-stimulus ratio *λ* = 0.0 for clarity.

To further probe the fast NMM’s gated oscillations, we computed the IEPC across a 0.5s window centred at the onset of each phoneme in the stimulus period (5.0 to 10.0s) of the 20 trials of the first ‘b’ condition item. Figure 9 **a** presents the IEPC across these 340 phoneme onsets and **b** presents the IEPC with the mean IEPC between −0.25 and −0.15s of each frequency band subtracted to show the transient response. There is a high IEPC at 4Hz and its harmonics due to the entrainment to the stimulus rate, alongside a high frequency peak at around 28.5Hz. This high frequency peak is revealed to be a transient response to the phoneme onset in figure 9 **b**, where the IEPC, with a baseline from before the onset subtracted, contains only high IEPC at high frequency post stimulus onset.

**Figure 9:**
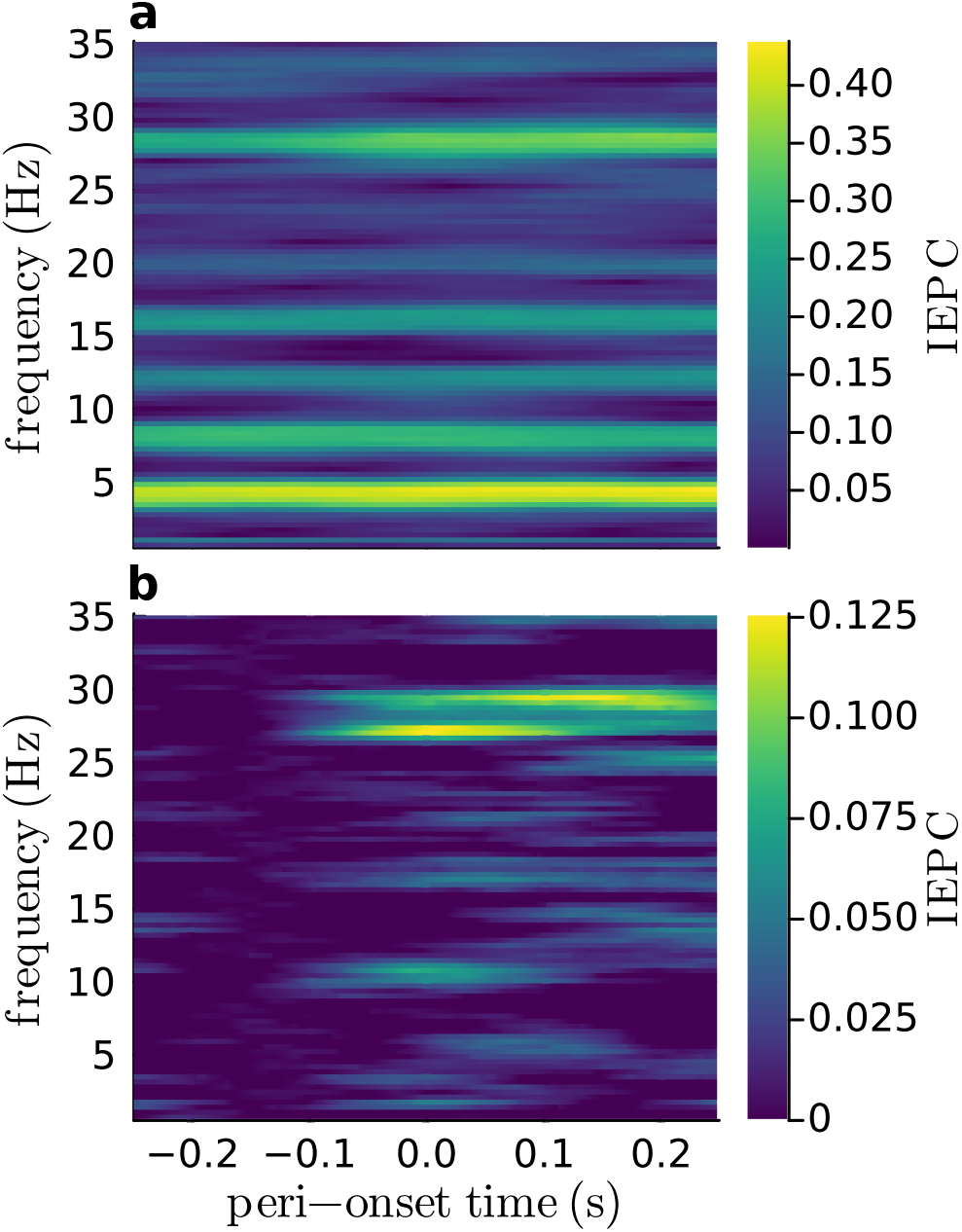
IEPC spectrograms showing transient phase-locking response to phoneme onsets. **a**: IEPC over all phoneme onsets of the “b” condition. **b**: IEPC minus the mean IEPC in the time range −0.25 to −0.15. There is strong 4Hz (and its harmonics) coherence across all phonemes due to the periodic nature of the stimulus. In addition to this, there is high phase coherence at around 30Hz caused by the phase reset fast oscillations. The baseline-removed plot illustrates that this is transient and locked to the onset of the phoneme. An exploration of the 4Hz IEPC after phoneme offset is also presented in the extended data fig. 9-1.

Finally, we conducted an exploratory analysis of the 4Hz IEPC for the ‘b’ syllable condition with the stimulus drive strength set to zero after ten seconds. This was to identify if the entrainment at the syllabic rate was sustained or transient post stimulus offset. We found that the 4Hz IEPC rapidly decreases after the end of the stimulus period (see extended data fig. 9-9). This indicates that the NMM only transiently entrains at the syllabic rate, similar to the experimental MEG findings of Oganian et al. (2023).

## Discussion

Our results show that the experimentally observed sharpness-specific tuning of neural entrainment to rhythmic speech can be reproduced using all three models considered here. The first model was a bio-physically-inspired neural mass model, the NMM. The other two models were phenomenological, representing the phase-reset and evoked-response mechanisms that have been proposed as sources of neural speech tracking. Although both NMM’s and both phenomenological models reproduced the correlation, they did so with differing stimulus requirements. The evoked model needed a pre-processed form of the stimulus, for instance, the peak-rate impulses. The NMM and phase-resetting oscillator, however, succeeded when processing just the continuous envelope of the syllables. While these models therefore appear to provide a more parsimonious explanation of sharpness-specific tuning of neural speech tracking, this contradicts the neurophysiological observation that auditory cortex predominantly represents acoustic edges (peak-rate events), rather than the continuous envelope (Oganian and Chang, 2019). As such, this result alone is not enough to decide between evoked or phase-resetting mechanisms, nor to conclude that the NMM and phase-resetting oscillator are better models of sharpness-specific tuning.

To resolve this ambiguity, we moved beyond just matching high-level correlations in the response of the models and explored their dynamics. In the spirit of Doelling and Assaneo (2021), we aim to identify specific biophysical mechanisms that may underpin these processes. Our tests probing the dynamics of the models provide insight into a set of behaviours that suggest that the behaviour of the NMM and phase-resetting oscillator are distinct and that the NMM is an excellent candidate for biophysical modelling of neural speech processing.

First, we found in the PCM test that both the slow and fast NMMs had phase-lags that varied as the stimulus frequency varied. This was in contrast to the concentrated phase-lags produced by the tuned phase-resetting oscillator. A shifting phase lag in the PCM test is indicative of evoked-like behaviour, which is somewhat surprising for the NMM given its oscillatory nature. While the tuned phase-resetting oscillator entrains to the gradual modulations of the sine wave, the NMM appears to exhibit a thresholding effect for phase-resetting that causes the evoked-like behaviour in this test. When driven by the slowly varying sine waves, the NMM does not phase reset, and is instead smoothly-modulated, resulting in the shifting phase lag. This suggests that the NMM can act like a linear filter for smoothly-varying signals. This is made evident in the syllabic rate test and subsequent investigation into the phase-resetting dynamics of the NMM.

The evidence for the thresholded phase-resetting dynamics comes from the different result in the syllabic rate test. Here the NMMs behave most like a phase-resetting oscillator. In particular, the slow NMM exhibits a peak in ITPC centred at its own natural frequency. This is similar to the sharp 4Hz ITPC peak of the 4Hz phase-resetting oscillator, but broader, indicating that the slow NMM can entrain to syllabic rates that diverge from its natural frequency. The fast NMM exhibits high ITPC across a broad range of frequencies that matches or outperforms the tuned phase-resetting oscillator within the tested frequency range. In this test, the models are driven by the CV-syllable envelopes that were used for the sharpness-specific tuning experiment. These contain broadband energy and sharp onsets that are sufficient to cause phase-resetting in the NMMs where the smooth narrowband sine-wave stimuli do not. Figures 7 and 8 demonstrate phase-resetting in both the slow and fast NMMs.

The fast NMM is able to track widely varying syllabic rates, similar to evoked models and the tuned phase-resetting oscillator. A possible explanation for this is the cross-frequency coupling between the slow syllabic rate and the fast intrinsic rhythm that is visible in fig. 7. This may allow even slow syllabic rates to strongly entrain the fast NMM as the sharp onsets of the phonemes trigger phase-reset high-frequency oscillations that are gated by the slow syllabic rate. Faster syllabic rates presumably may just entrain the baseline oscillation of the model, resulting in the broad peak of ITPC for the fast NMM. Recent views have stressed the importance of using non-isochronous stimuli when evaluating the algorithms of neural speech tracking to not unjustifiably favour oscillatory mechanisms that rely upon resonance at a fixed frequency (Lalor and Nidiffer, 2025). Although our syllabic rate test used effectively isochronus stimuli, the fast NMM’s robust entrainment across rates suggests that it may also be able to entrain to natural speech’s variable rate. This is in contrast to the simple phase-resetting oscillator, where the rapid loss of entrainment at rates distinct from 4Hz demonstrates its dependence upon the rhythmic nature of our stimuli.

This cross-frequency coupling has in fact been used as a test to differentiate between the mechanisms underlying speech tracking. Oganian et al. (2023) found that when listening to slowed natural speech, a peak in IEPC was present post syllable onset at the mean syllabic rate (1.9Hz) alongside a peak in the Theta band. By modelling this with a linear evoked model and a phase-resetting oscillator, they demonstrated that only the evoked model could reproduce this dual peak, attributing the low frequency IEPC peak to the syllabic rate, and the high frequency peak to the frequency content of the evoked response itself. Here, we inspected the IEPC of the fast NMM’s response to our stimuli to explore the high frequency nested oscillations further. We found that the fast NMM’s IEPC has features in common with the dual spectral peak observed by Oganian et al. (2023): Strong entrainment at the stimulus frequency (4Hz), and transient entrainment at a higher frequency. This qualitatively demonstrates that an alternate mechanism could be responsible for this feature. Namely, a nested high frequency oscillation that is transiently phase-reset by syllable onsets. And further, that this can be provided natively by a mean-field approximation of a biophysical network of spiking neurons; the NMM. The NMM’s response may also address recent concerns about the compatibility of generative oscillatory mechanisms with temporal response functions derived from the continuous envelope (Lalor and Nidiffer, 2025). Our results demonstrate how a nonlinear oscillatory process can respond to sharp onsets in the continuous envelope, producing nested oscillations that are free to play out between onsets.

The aim of exploring the use of the NMM is to see if we can obtain more biophysical understanding of the mechanisms responsible for speech processing in the brain. As mentioned above, an interesting biophysical fact about speech tracking is that activity in the STG represents peak-rate events rather than the continuous envelope of the speech (Oganian and Chang, 2019). We suggest that the NMM provides a mechanistic bridge between the physiology and these observations. The NMM’s activity represents the behaviour of a large population of spiking neurons. We have demonstrated that when driven by the continuous envelope, these neurons effectively detect sharp onsets in the envelope, phase-locking to them. Further, it is doing so while demonstrating experimentally validated hallmarks of this processing, such as the dual-peak IEPC signature and sharpness-specific tuning of the entrainment. The peak-rate event features identified in STG activity may therefore be the result of non-linear dynamics such as those of the NMM.

While these results provide a mechanistic candidate for modelling neural speech tracking, our investigation has several limitations. Most notably, we only qualitatively matched the dual-peak IEPC result from Oganian et al. (2023). The peaks observed in MEG occurred specifically at the stimulus rate and in the Theta band. We instead have a peak at high-beta/low-gamma, at around 30Hz. This is a significant difference given the functionally distinct roles of theta and beta/gamma rhythms. Indeed, beta oscillations are typically suppressed during speech production and motor tasks. However, this discrepancy is due to the parameters of the fast NMM being derived to model motor beta-rhythms (Byrne et al., 2017). This parameter set was used to explore whether the NMM would entrain when its un-perturbed frequency was distinct from the 4Hz syllabic rate. The emergence of this qualitative match to the dual-peak IEPC feature is notable, but these parameters should be tuned to determine whether a quantitative match is achievable. In addition, our test used the near-isochronous CV-syllable stimuli from the experiment by Cucu et al. (2022). This is a known confound when trying to evaluate oscillatory and evoked tracking (Lalor and Nidiffer, 2025) that partially obscured the transient phase-locking response to phoneme onsets in the IEPC test with persistent entrainment to the stimulus rate both pre- and post-onset. Further, our exploration of the simulation noise level and drive strength was quite coarse. Sensitivity analysis at a finer resolution could explore how robust the functional operating point is.

There are many opportunities for further computational investigations to confirm these results and to explore whether the NMM can support other features of neural speech processing. For instance, the NMM’s cross-frequency coupling could be tested using natural, temporally jittered speech to isolate the transient phase-resetting behaviour without rhythmic interference. As was carried out in Oganian et al. (2023), the wide distribution of inter-syllable timings in natural speech could also be used to investigate the absence of sustained syllabic-rate entrainment more rigorously than was possible through our inspection of the end of our periodic stimulus. Further, the cross-frequency coupling could be explored to see if it may correspond to the proposed hierarchical oscillatory structure suggested by Giraud and Poeppel (2012). Multi-region architectures may also be considered, potentially offering a way to capture the balance between bottom-up sensory processing and top-down attentional modulation in speech processing (Poeppel and Assaneo, 2020). They may also produce a better background noise representation than the stimulus-based noise used here. Adding stochasticity to the dynamics beyond the noise added to the stimulus and post-hoc 1*/f* noise could result in more realistic results. Further study of the phase-reset and evoked response models could use filter-bank based models to represent tonotopic brain structures, alongside interrogation of other parameter sensitivities such as the time scale of the evoked-response model. The biological plausibility of these results could be strengthened by exploring parameter ranges that are restricted to experimentally validated ranges. Subsequent tuning of these parameters could result in a closer match to experimental results. For example, tuning the synaptic timescales or membrane time constants of the NMM may shift the high IEPC peak down towards the theta range.

To confirm the biophysical mechanisms proposed by these models, targeted experimental work is required. For example, the divergence of particular phoneme groups from the experimental trend order suggests that spectral content may play a role in the final entrainment observed. An experiment following the same procedure as Cucu et al. (2022)’s with the phonemes distorted to maintain constant sharpness across conditions may reveal the effect of the spectral content of the sounds. Additionally, the correlation of phoneme sharpness with syllabic rate tracking could be investigated under natural speech conditions. This would also enable an evaluation of the NMM’s transient IEPC response to natural speech.

## Supporting information

Manuscript .tex file and figures (.eps).

## Conflict of interest statement

The authors declare no competing interests

## Acknowledgements

We are grateful to Oana Cucu for the provision of EEG data and experimental stimuli that were central to this work, and Mark Toolan for preliminary work performed as part of his master’s thesis. AS is funded by a UK Research and Innovation grant (EP/S022937/1). This work was carried out using the computational facilities of the Advanced Computing Research Centre, University of Bristol - http://www.bris.ac.uk/acrc/. For the purpose of open access, the authors have applied a Creative Commons Attribution (CC BY) licence to any Author Accepted Manuscript version arising from this sub-mission.

## Extended Data

**Figure 1-1:**
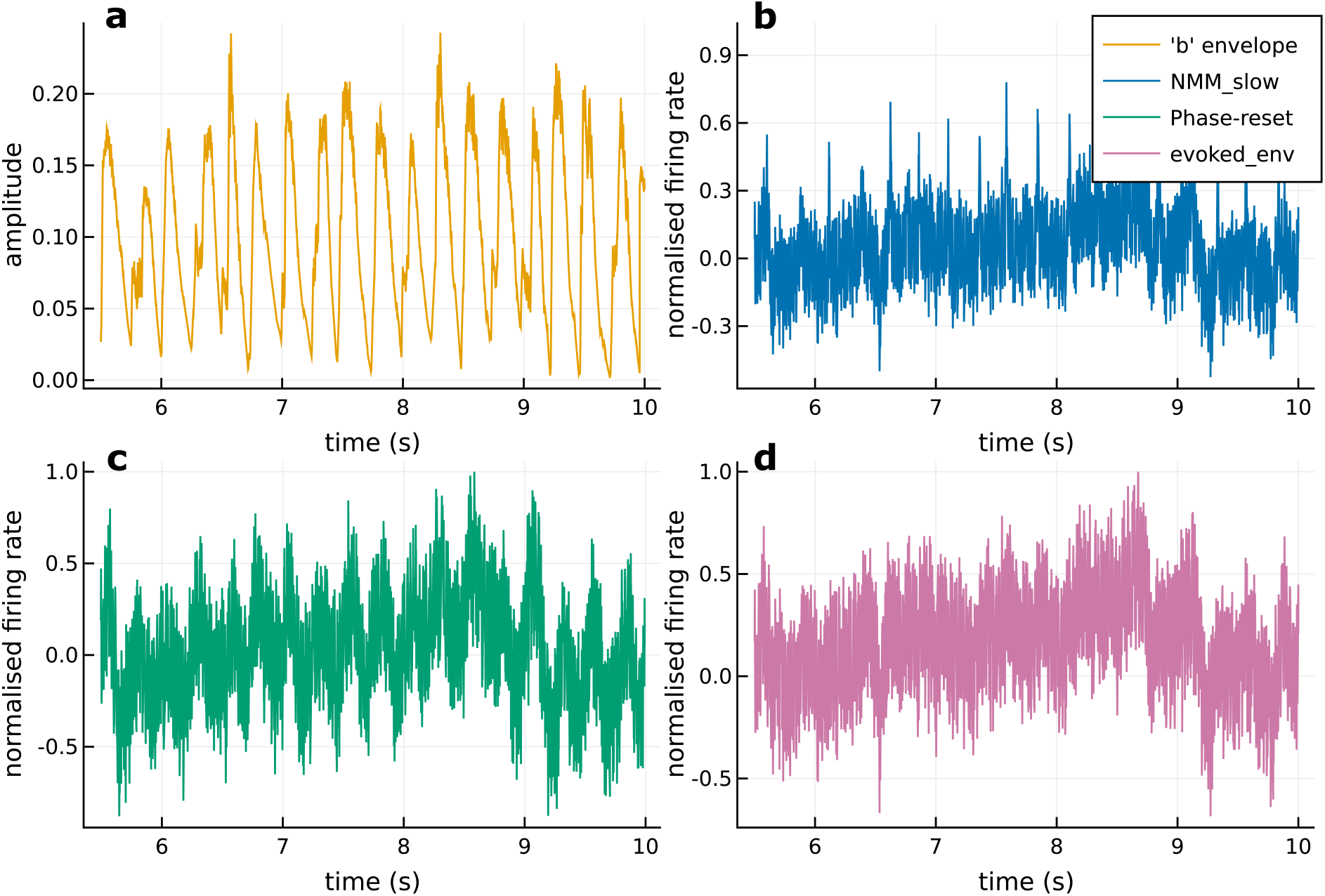
Example of the response of each model to a given consonant-vowel syllable sequence stimulus. **a**: Envelope from one of the “b” items with noise, *λ* = 0.7. **b**: Response of the slow NMM variant. **c**: Response of the phase-resetting oscillator. **d**: Response of the evoked-response model. All responses include post-hoc noise added at a signal to noise ratio of 0.5

**Figure 3-1:**
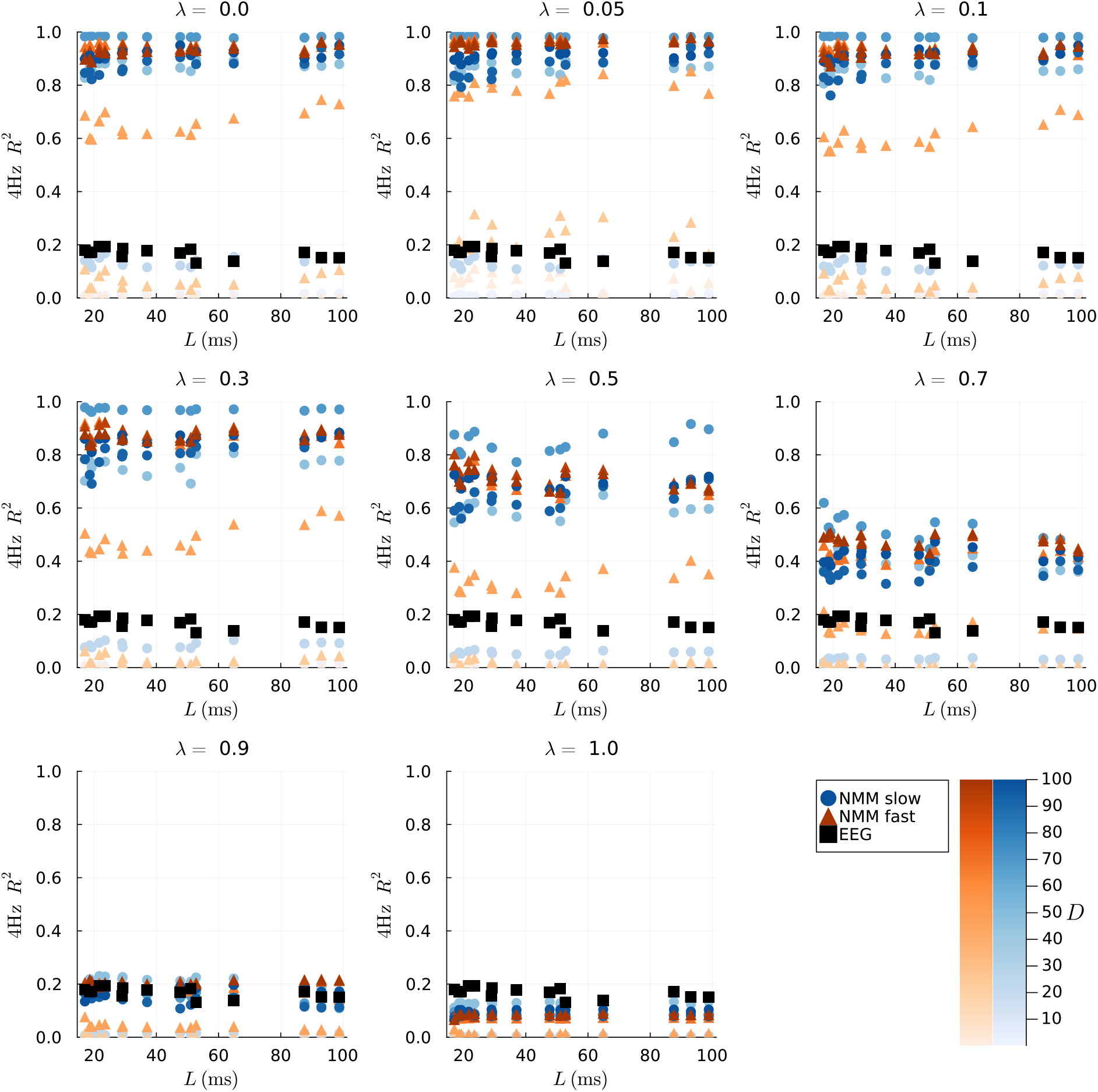
Mean 4Hz *R*^2^ across each phoneme condition’s three items vs *L* for both the slow (blue) and fast (orange) NMM models. Each subplot presents the results across values of *D*, with the colour intensity increasing with increasing *D*, for a particular *λ*. EEG result from Cucu et al. (2022) included for comparison. Higher *D* generally increases the entrainment strength. A sufficient level of noise *λ* is required to obtain an entrainment strength comparable to EEG experiment.

**Figure 3-2:**
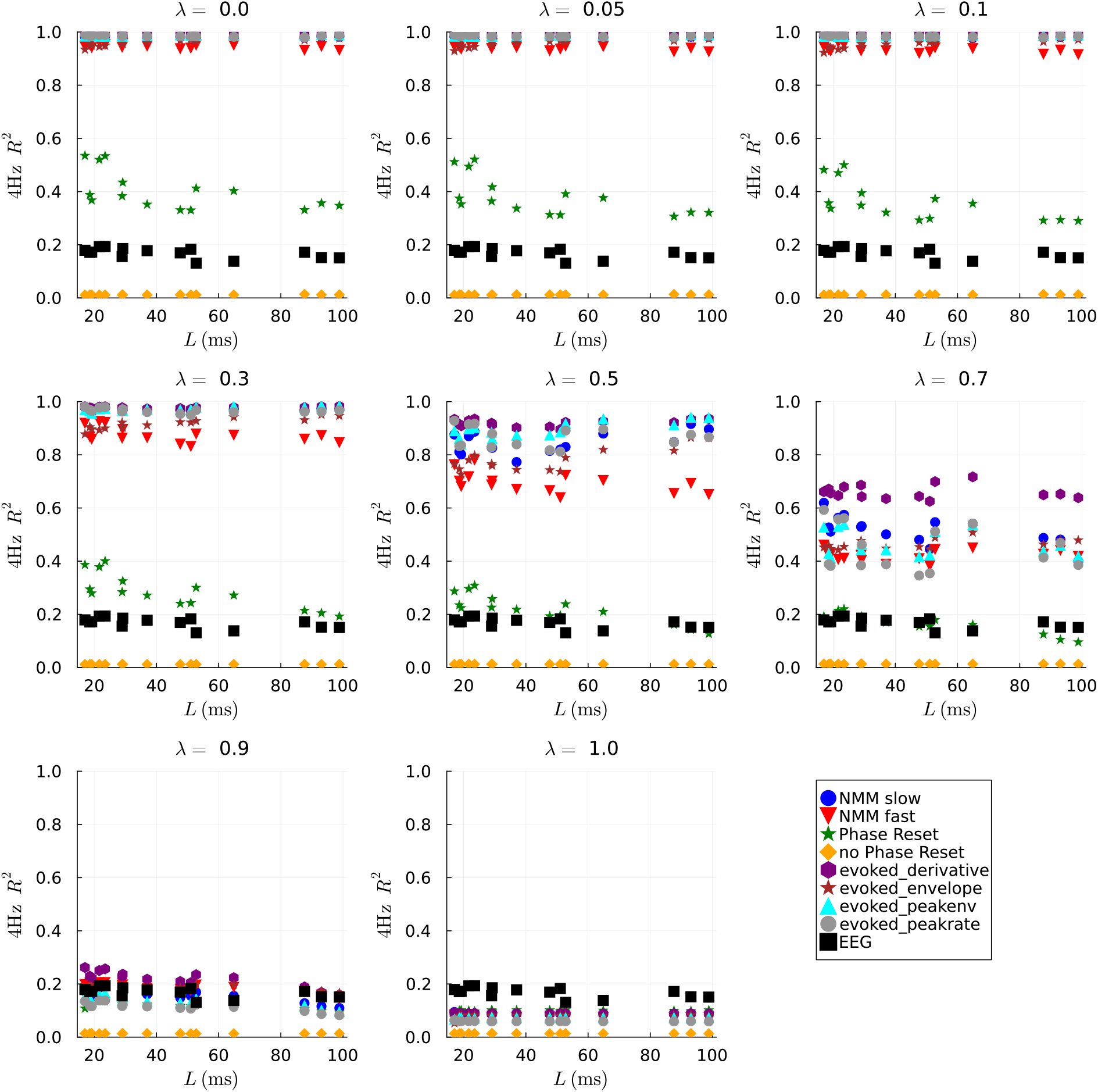
Mean 4Hz *R*^2^ across each phoneme condition’s three items vs *L* for all of the models tested in this work. Each subplot presents the results for a particular *λ*. NMM results are for *D* = 12.2. EEG result from Cucu et al. (2022) included for comparison.

**Figure 9-1:**
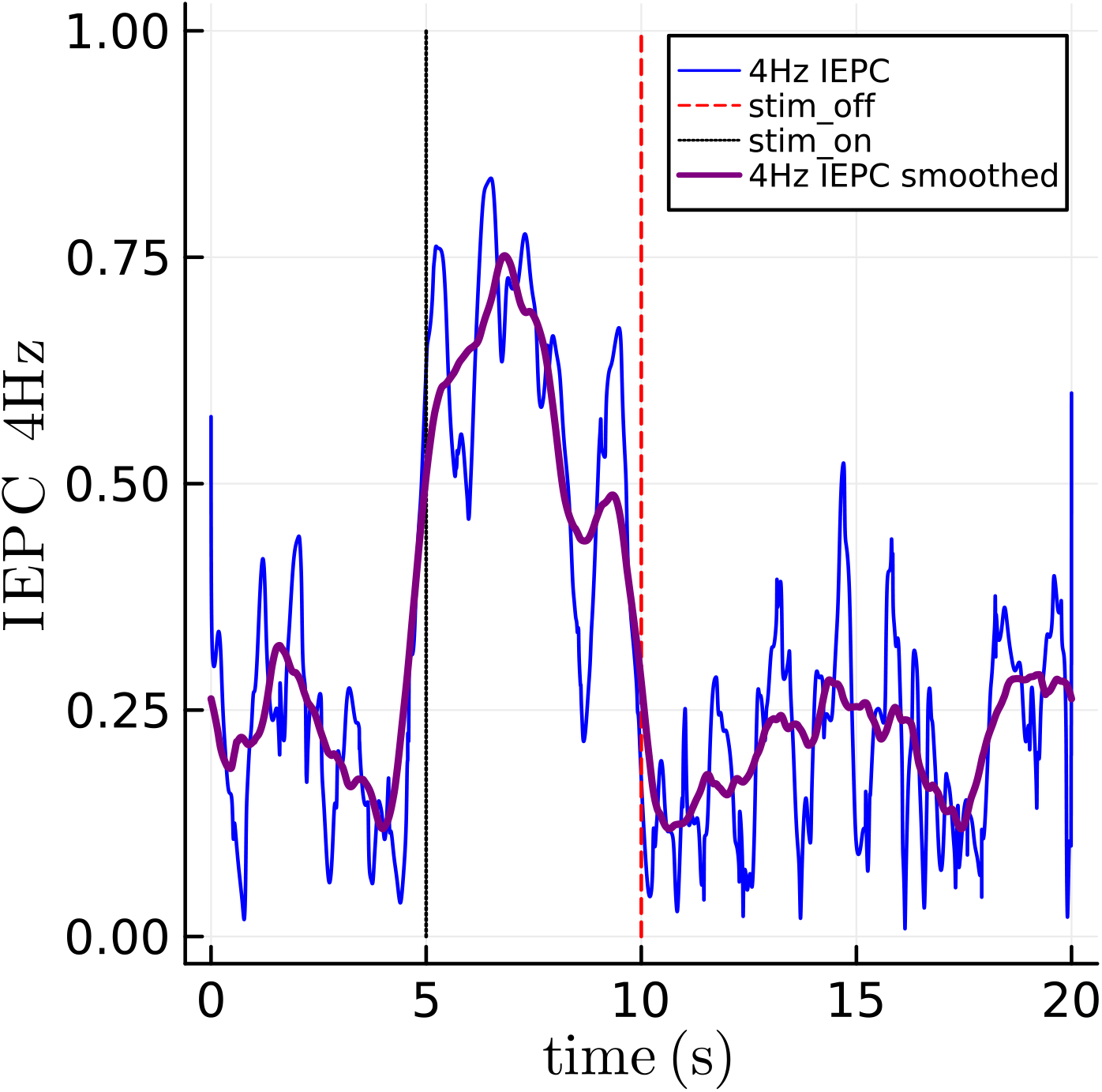
4Hz IEPC across the 60 ‘b’ condition trials. The first 5.0 seconds are the noisy syllable based drive, which changes to the 4Hz drive with *λ* = 0 from 5.0 to 10.0s. From 10.0 seconds onwards, the drive strength is set to zero. Blue line, the raw 4Hz IEPC. Black dotted line, the isochronous stimulus onset point. Red dashed line, the stimulus offset point. Purple thicker line, a 1.2s window zero-phase moving average smoothed version of the 4Hz IEPC. Once the isochronous stimulus starts, the NMM is entrained to the stimulus. At cessation, the entrainment drops back to the noisy baseline level indicating no ringing in the entrainment at the syllabic rate post stimulus offset.

**Table 3-1:** Correlation between phoneme sharpness and 4Hz ITPC for the fast NMM variant across *λ* and *D* values. We present uncorrected and corrected *p* values for the correlations. Bonferroni and Benjamini-Hochberg FDR corrections were calculated for eight comparisons (the model and stimulus variants) and a false discovery rate of 0.05.

| Model | $\lambda$ | $D$ | $r$ | CI<sub>lower</sub> | CI<sub>upper</sub> | $p$ | $p_{\text{Bonferroni}}$ | $p_{\text{FDR}}$ |
| --- | --- | --- | --- | --- | --- | --- | --- | --- |
| NMM fast | 0 | 0.1 | -0.852 | -0.95 | -0.604 | 5.417e-05 | 0.000433 | 0.000157 |
| NMM fast | 0 | 0.496 | 0.269 | -0.282 | 0.687 | 0.332 | 1 | 0.379 |
| NMM fast | 0 | 2.46 | 0.66 | 0.222 | 0.876 | 0.00748 | 0.0598 | 0.0199 |
| NMM fast | 0 | 12.2 | -0.585 | -0.844 | -0.104 | 0.022 | 0.176 | 0.0293 |
| NMM fast | 0 | 60.5 | 0.632 | 0.177 | 0.865 | 0.0114 | 0.0916 | 0.0183 |
| NMM fast | 0 | 100 | 0.764 | 0.414 | 0.917 | 0.000912 | 0.0073 | 0.00243 |
| NMM fast | 0.05 | 0.1 | -0.477 | -0.795 | 0.0469 | 0.0723 | 0.579 | 0.116 |
| NMM fast | 0.05 | 0.496 | 0.129 | -0.41 | 0.602 | 0.647 | 1 | 0.647 |
| NMM fast | 0.05 | 2.46 | 0.437 | -0.097 | 0.776 | 0.103 | 0.827 | 0.207 |
| NMM fast | 0.05 | 12.2 | -0.144 | -0.611 | 0.397 | 0.608 | 1 | 0.608 |
| NMM fast | 0.05 | 60.5 | 0.517 | 0.00638 | 0.814 | 0.0485 | 0.388 | 0.0969 |
| NMM fast | 0.05 | 100 | 0.575 | 0.089 | 0.84 | 0.0249 | 0.199 | 0.0665 |
| NMM fast | 0.1 | 0.1 | -0.852 | -0.95 | -0.604 | 5.422e-05 | 0.000434 | 0.000175 |
| NMM fast | 0.1 | 0.496 | 0.177 | -0.369 | 0.632 | 0.529 | 1 | 0.529 |
| NMM fast | 0.1 | 2.46 | 0.832 | 0.557 | 0.942 | 0.000121 | 0.000966 | 0.000322 |
| NMM fast | 0.1 | 12.2 | -0.588 | -0.845 | -0.108 | 0.0212 | 0.17 | 0.0283 |
| NMM fast | 0.1 | 60.5 | 0.439 | -0.0943 | 0.777 | 0.101 | 0.812 | 0.116 |
| NMM fast | 0.1 | 100 | 0.725 | 0.339 | 0.902 | 0.00221 | 0.0177 | 0.00441 |
| NMM fast | 0.3 | 0.1 | -0.844 | -0.947 | -0.586 | 7.472e-05 | 0.000598 | 0.000299 |
| NMM fast | 0.3 | 0.496 | 0.0161 | -0.5 | 0.524 | 0.954 | 1 | 0.954 |
| NMM fast | 0.3 | 2.46 | 0.828 | 0.549 | 0.941 | 0.000136 | 0.00109 | 0.000545 |
| NMM fast | 0.3 | 12.2 | -0.479 | -0.796 | 0.0441 | 0.0709 | 0.567 | 0.113 |
| NMM fast | 0.3 | 60.5 | -0.0262 | -0.531 | 0.493 | 0.926 | 1 | 0.926 |
| NMM fast | 0.3 | 100 | 0.39 | -0.152 | 0.752 | 0.15 | 1 | 0.172 |
| NMM fast | 0.5 | 0.1 | -0.818 | -0.937 | -0.526 | 0.000194 | 0.00155 | 0.000777 |
| NMM fast | 0.5 | 0.496 | -0.186 | -0.637 | 0.361 | 0.508 | 1 | 0.58 |
| NMM fast | 0.5 | 2.46 | 0.488 | -0.0322 | 0.8 | 0.0649 | 0.519 | 0.13 |
| NMM fast | 0.5 | 12.2 | -0.517 | -0.814 | -0.0064 | 0.0485 | 0.388 | 0.0969 |
| NMM fast | 0.5 | 60.5 | -0.634 | -0.865 | -0.18 | 0.0112 | 0.0896 | 0.0179 |
| NMM fast | 0.5 | 100 | -0.522 | -0.816 | -0.0135 | 0.0459 | 0.367 | 0.0917 |
| NMM fast | 0.7 | 0.1 | -0.751 | -0.912 | -0.388 | 0.00125 | 0.00997 | 0.00499 |
| NMM fast | 0.7 | 0.496 | -0.329 | -0.72 | 0.221 | 0.231 | 1 | 0.617 |
| NMM fast | 0.7 | 2.46 | -0.0897 | -0.576 | 0.443 | 0.751 | 1 | 0.751 |
| NMM fast | 0.7 | 12.2 | 0.147 | -0.395 | 0.613 | 0.602 | 1 | 0.705 |
| NMM fast | 0.7 | 60.5 | -0.178 | -0.633 | 0.368 | 0.525 | 1 | 0.7 |
| NMM fast | 0.7 | 100 | -0.477 | -0.795 | 0.0471 | 0.0724 | 0.579 | 0.193 |
| NMM fast | 0.9 | 0.1 | -0.57 | -0.838 | -0.0814 | 0.0265 | 0.212 | 0.0354 |
| NMM fast | 0.9 | 0.496 | -0.306 | -0.707 | 0.245 | 0.267 | 1 | 0.318 |
| NMM fast | 0.9 | 2.46 | -0.709 | -0.896 | -0.309 | 0.00307 | 0.0245 | 0.00491 |
| NMM fast | 0.9 | 12.2 | -0.909 | -0.97 | -0.742 | 2.716e-06 | 2.173e-05 | 2.173e-05 |
| NMM fast | 0.9 | 60.5 | 0.767 | 0.42 | 0.918 | 0.000844 | 0.00675 | 0.00113 |
| NMM fast | 0.9 | 100 | 0.564 | 0.0729 | 0.835 | 0.0285 | 0.228 | 0.0456 |
| NMM fast | 1 | 0.1 | -0.338 | -0.725 | 0.211 | 0.218 | 1 | 0.34 |
| NMM fast | 1 | 0.496 | -0.28 | -0.693 | 0.271 | 0.312 | 1 | 0.357 |
| NMM fast | 1 | 2.46 | -0.156 | -0.619 | 0.387 | 0.579 | 1 | 0.579 |
| NMM fast | 1 | 12.2 | -0.183 | -0.636 | 0.363 | 0.514 | 1 | 0.514 |
| NMM fast | 1 | 60.5 | 0.509 | -0.00474 | 0.81 | 0.0528 | 0.422 | 0.249 |
| NMM fast | 1 | 100 | 0.208 | -0.34 | 0.651 | 0.456 | 1 | 0.456 |

**Table 3-2:**
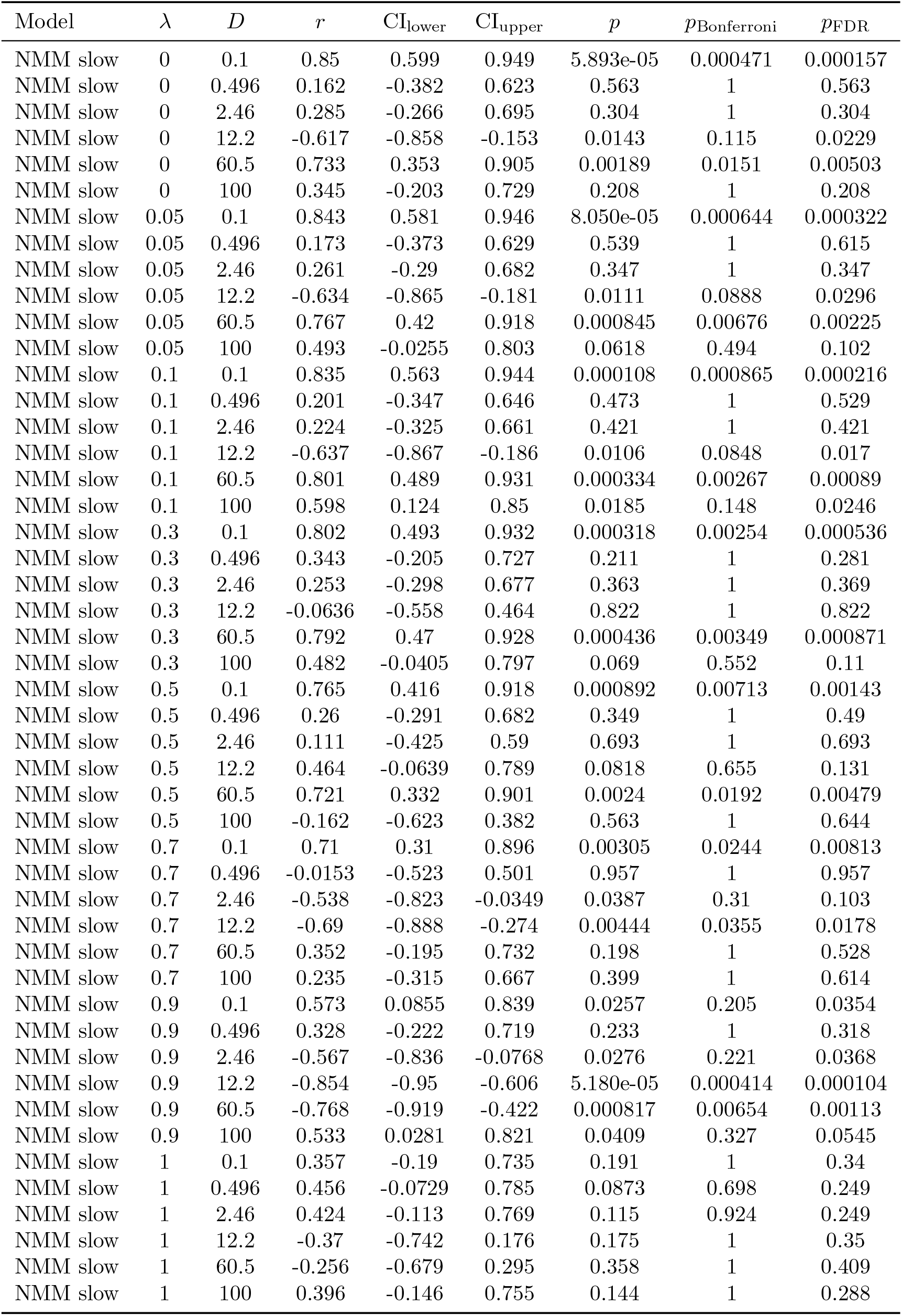
Correlation between phoneme sharpness and 4Hz ITPC for the slow NMM variant across *λ* and *D* values. We present uncorrected and corrected *p* values for the correlations. Bonferroni and Benjamini-Hochberg FDR corrections were calculated for eight comparisons (the model and stimulus variants) and a false discovery rate of 0.05.

| Model | $\lambda$ | $D$ | $r$ | $CI_{\text{lower}}$ | $CI_{\text{upper}}$ | $p$ | $p_{\text{Bonferroni}}$ | $p_{\text{FDR}}$ |
| --- | --- | --- | --- | --- | --- | --- | --- | --- |
| NMM slow | 0 | 0.1 | 0.85 | 0.599 | 0.949 | 5.893e-05 | 0.000471 | 0.000157 |
| NMM slow | 0 | 0.496 | 0.162 | -0.382 | 0.623 | 0.563 | 1 | 0.563 |
| NMM slow | 0 | 2.46 | 0.285 | -0.266 | 0.695 | 0.304 | 1 | 0.304 |
| NMM slow | 0 | 12.2 | -0.617 | -0.858 | -0.153 | 0.0143 | 0.115 | 0.0229 |
| NMM slow | 0 | 60.5 | 0.733 | 0.353 | 0.905 | 0.00189 | 0.0151 | 0.00503 |
| NMM slow | 0 | 100 | 0.345 | -0.203 | 0.729 | 0.208 | 1 | 0.208 |
| NMM slow | 0.05 | 0.1 | 0.843 | 0.581 | 0.946 | 8.050e-05 | 0.000644 | 0.000322 |
| NMM slow | 0.05 | 0.496 | 0.173 | -0.373 | 0.629 | 0.539 | 1 | 0.615 |
| NMM slow | 0.05 | 2.46 | 0.261 | -0.29 | 0.682 | 0.347 | 1 | 0.347 |
| NMM slow | 0.05 | 12.2 | -0.634 | -0.865 | -0.181 | 0.0111 | 0.0888 | 0.0296 |
| NMM slow | 0.05 | 60.5 | 0.767 | 0.42 | 0.918 | 0.000845 | 0.00676 | 0.00225 |
| NMM slow | 0.05 | 100 | 0.493 | -0.0255 | 0.803 | 0.0618 | 0.494 | 0.102 |
| NMM slow | 0.1 | 0.1 | 0.835 | 0.563 | 0.944 | 0.000108 | 0.000865 | 0.000216 |
| NMM slow | 0.1 | 0.496 | 0.201 | -0.347 | 0.646 | 0.473 | 1 | 0.529 |
| NMM slow | 0.1 | 2.46 | 0.224 | -0.325 | 0.661 | 0.421 | 1 | 0.421 |
| NMM slow | 0.1 | 12.2 | -0.637 | -0.867 | -0.186 | 0.0106 | 0.0848 | 0.017 |
| NMM slow | 0.1 | 60.5 | 0.801 | 0.489 | 0.931 | 0.000334 | 0.00267 | 0.00089 |
| NMM slow | 0.1 | 100 | 0.598 | 0.124 | 0.85 | 0.0185 | 0.148 | 0.0246 |
| NMM slow | 0.3 | 0.1 | 0.802 | 0.493 | 0.932 | 0.000318 | 0.00254 | 0.000536 |
| NMM slow | 0.3 | 0.496 | 0.343 | -0.205 | 0.727 | 0.211 | 1 | 0.281 |
| NMM slow | 0.3 | 2.46 | 0.253 | -0.298 | 0.677 | 0.363 | 1 | 0.369 |
| NMM slow | 0.3 | 12.2 | -0.0636 | -0.558 | 0.464 | 0.822 | 1 | 0.822 |
| NMM slow | 0.3 | 60.5 | 0.792 | 0.47 | 0.928 | 0.000436 | 0.00349 | 0.000871 |
| NMM slow | 0.3 | 100 | 0.482 | -0.0405 | 0.797 | 0.069 | 0.552 | 0.11 |
| NMM slow | 0.5 | 0.1 | 0.765 | 0.416 | 0.918 | 0.000892 | 0.00713 | 0.00143 |
| NMM slow | 0.5 | 0.496 | 0.26 | -0.291 | 0.682 | 0.349 | 1 | 0.49 |
| NMM slow | 0.5 | 2.46 | 0.111 | -0.425 | 0.59 | 0.693 | 1 | 0.693 |
| NMM slow | 0.5 | 12.2 | 0.464 | -0.0639 | 0.789 | 0.0818 | 0.655 | 0.131 |
| NMM slow | 0.5 | 60.5 | 0.721 | 0.332 | 0.901 | 0.0024 | 0.0192 | 0.00479 |
| NMM slow | 0.5 | 100 | -0.162 | -0.623 | 0.382 | 0.563 | 1 | 0.644 |
| NMM slow | 0.7 | 0.1 | 0.71 | 0.31 | 0.896 | 0.00305 | 0.0244 | 0.00813 |
| NMM slow | 0.7 | 0.496 | -0.0153 | -0.523 | 0.501 | 0.957 | 1 | 0.957 |
| NMM slow | 0.7 | 2.46 | -0.538 | -0.823 | -0.0349 | 0.0387 | 0.31 | 0.103 |
| NMM slow | 0.7 | 12.2 | -0.69 | -0.888 | -0.274 | 0.00444 | 0.0355 | 0.0178 |
| NMM slow | 0.7 | 60.5 | 0.352 | -0.195 | 0.732 | 0.198 | 1 | 0.528 |
| NMM slow | 0.7 | 100 | 0.235 | -0.315 | 0.667 | 0.399 | 1 | 0.614 |
| NMM slow | 0.9 | 0.1 | 0.573 | 0.0855 | 0.839 | 0.0257 | 0.205 | 0.0354 |
| NMM slow | 0.9 | 0.496 | 0.328 | -0.222 | 0.719 | 0.233 | 1 | 0.318 |
| NMM slow | 0.9 | 2.46 | -0.567 | -0.836 | -0.0768 | 0.0276 | 0.221 | 0.0368 |
| NMM slow | 0.9 | 12.2 | -0.854 | -0.95 | -0.606 | 5.180e-05 | 0.000414 | 0.000104 |
| NMM slow | 0.9 | 60.5 | -0.768 | -0.919 | -0.422 | 0.000817 | 0.00654 | 0.00113 |
| NMM slow | 0.9 | 100 | 0.533 | 0.0281 | 0.821 | 0.0409 | 0.327 | 0.0545 |
| NMM slow | 1 | 0.1 | 0.357 | -0.19 | 0.735 | 0.191 | 1 | 0.34 |
| NMM slow | 1 | 0.496 | 0.456 | -0.0729 | 0.785 | 0.0873 | 0.698 | 0.249 |
| NMM slow | 1 | 2.46 | 0.424 | -0.113 | 0.769 | 0.115 | 0.924 | 0.249 |
| NMM slow | 1 | 12.2 | -0.37 | -0.742 | 0.176 | 0.175 | 1 | 0.35 |
| NMM slow | 1 | 60.5 | -0.256 | -0.679 | 0.295 | 0.358 | 1 | 0.409 |
| NMM slow | 1 | 100 | 0.396 | -0.146 | 0.755 | 0.144 | 1 | 0.288 |

**Table 3-3:**
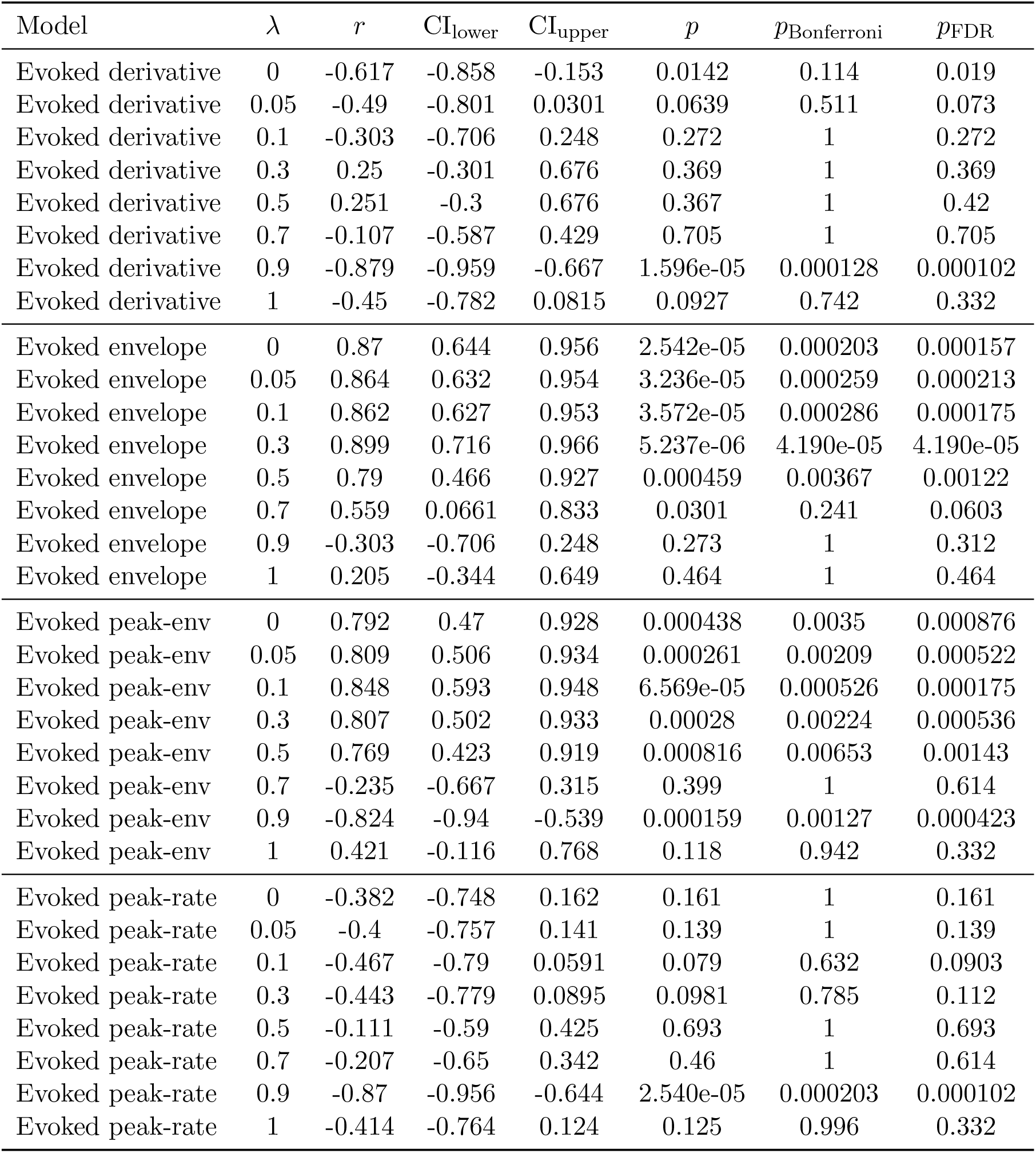
Correlation between phoneme sharpness and 4Hz ITPC for the evoked-response model for each stimulus variant across *λ* values. We present uncorrected and corrected *p* values for the correlations. Bonferroni and Benjamini-Hochberg FDR corrections were calculated for eight comparisons (the model and stimulus variants) and a false discovery rate of 0.05.

**Table 3-4:** Correlation between phoneme sharpness and 4Hz ITPC for both the phase-resetting and non phase-resetting oscillators across *λ* values. We present uncorrected and corrected *p* values for the correlations. Bonferroni and Benjamini-Hochberg FDR corrections were calculated for eight comparisons (the model and stimulus variants) and a false discovery rate of 0.05.

| Model | $\lambda$ | $r$ | CI<sub>lower</sub> | CI<sub>upper</sub> | $p$ | $p_{\text{Bonferroni}}$ | $p_{\text{FDR}}$ |
| --- | --- | --- | --- | --- | --- | --- | --- |
| No phase reset | 0 | 0.634 | 0.18 | 0.865 | 0.0112 | 0.0897 | 0.0179 |
| No phase reset | 0.05 | 0.675 | 0.249 | 0.882 | 0.00572 | 0.0458 | 0.00916 |
| No phase reset | 0.1 | 0.726 | 0.34 | 0.902 | 0.0022 | 0.0176 | 0.00351 |
| No phase reset | 0.3 | 0.513 | 0.0012 | 0.812 | 0.0504 | 0.403 | 0.0672 |
| No phase reset | 0.5 | 0.357 | -0.19 | 0.735 | 0.192 | 1 | 0.256 |
| No phase reset | 0.7 | 0.128 | -0.412 | 0.601 | 0.65 | 1 | 0.705 |
| No phase reset | 0.9 | 0.165 | -0.379 | 0.625 | 0.557 | 1 | 0.557 |
| No phase reset | 1 | -0.3 | -0.704 | 0.251 | 0.277 | 1 | 0.34 |
| Phase reset | 0 | -0.571 | -0.838 | -0.0837 | 0.026 | 0.208 | 0.0298 |
| Phase reset | 0.05 | -0.624 | -0.861 | -0.164 | 0.013 | 0.104 | 0.0173 |
| Phase reset | 0.1 | -0.658 | -0.875 | -0.219 | 0.00771 | 0.0617 | 0.0103 |
| Phase reset | 0.3 | -0.801 | -0.931 | -0.489 | 0.000335 | 0.00268 | 0.000536 |
| Phase reset | 0.5 | -0.865 | -0.954 | -0.634 | 3.099e-05 | 0.000248 | 0.000248 |
| Phase reset | 0.7 | -0.892 | -0.964 | -0.699 | 7.880e-06 | 6.304e-05 | 6.304e-05 |
| Phase reset | 0.9 | -0.785 | -0.925 | -0.457 | 0.00052 | 0.00416 | 0.00104 |
| Phase reset | 1 | 0.288 | -0.263 | 0.697 | 0.298 | 1 | 0.34 |

## Notes

https://github.com/Modelling-Speech-Processing/Shannon_et_al

### Competing Interest Statement

The authors have declared no competing interest.

### Summary of Updates

The manuscript has been extended with new tests exploring the dynamics of the NMM and how they compare to the phenomenological models' behaviour alongside extensive revision of the prose and main discussion points to highlight the new findings. New figures include Figures 3, 5, 6, 7, 8, and 9 and Figure 9-1 in the extended data.

